# Vaccinia E5 is a major inhibitor of the DNA sensor cGAS

**DOI:** 10.1101/2021.10.25.465197

**Authors:** Ning Yang, Yi Wang, Peihong Dai, Tuo Li, Christian Zierhut, Adrian Tan, Tuo Zhang, Heng Pan, Zhuoning Li, Alban Ordureau, Jenny Zhaoying Xiang, Ronald C. Hendrickson, Hironori Funabiki, Zhijian Chen, Liang Deng

## Abstract

The DNA sensor cyclic GMP-AMP synthase (cGAS) is critical in host antiviral immunity. Vaccinia virus (VACV) is a large cytoplasmic DNA virus that belongs to the poxvirus family. How vaccinia virus antagonizes the cGAS-mediated cytosolic DNA-sensing pathway is largely unknown. In this study, we screened 82 vaccinia viral genes to identify potential viral inhibitors of the cGAS/Stimulator of interferon gene (STING) pathway. We discovered that vaccinia E5 is a virulence factor and a major inhibitor of cGAS that elicits proteasome-dependent cGAS degradation. E5 localizes to the cytoplasm and nuclei of infected cells. Cytosolic E5 triggers K48-linked ubiquitination of cGAS and proteasome-dependent degradation via interacting with cGAS. E5 itself also undergoes ubiquitination and degradation. Deleting the E5R gene from the Modified vaccinia virus Ankara (MVA) genome strongly induces type I IFN production by dendritic cells (DCs) and promotes DC maturation, thereby improving the immunogenicity of the viral vector.

## INTRODUCTION

Cyclic GMP-AMP synthase (cGAS) is a major cytosolic DNA sensor critical to antiviral, antitumor innate immunity, as well as in autoimmune inflammatory diseases (Ablasser and Chen, 2019; Li et al., 2013; Schoggins et al., 2014; Wu et al., 2013). Once activated by cytosolic DNA, cGAS generates cyclic GMP-AMP (cGAMP), which in turn binds to an endoplasmic reticulum-localized protein STING, resulting in the activation of the TBK1/IRF3/IFNB pathway. Consequently, viruses have evolved to employ many strategies to evade this important antiviral pathway (Lau et al., 2015; Ma and Damania, 2016; Wu et al., 2015; Zhang et al., 2016).

Poxviruses are large cytoplasmic DNA viruses that are important human and veterinary pathogens as well as oncolytic agents and viral vectors. Vaccinia virus (VACV) was used successfully as a vaccine for smallpox eradication. However, direct infection of dendritic cells (DCs) with vaccinia results in inhibition of both innate and adaptive immune responses (Deng et al., 2006; Engelmayer et al., 1999; Jenne et al., 2000). Modified vaccinia virus Ankara (MVA) is a highly attenuated vaccinia strain with deletion of large fragments from its parental vaccinia genome following more than 570 serial passages in chicken embryo fibroblasts, rendering it non-replicative in most mammalian cells (Antoine et al., 1998; Sutter and Moss, 1992). MVA is an important vaccine vector and was recently approved as a second-generation vaccine against smallpox and monkeypox (Pittman et al., 2019; Volz and Sutter, 2017). Unlike wild-type VACV, MVA infection of bone marrow-derived dendritic cells induces type I IFN in a cGAS/STING-dependent manner (Dai et al., 2014).

cGAS is important for host defense against poxvirus infection. cGAS-deficient mice are more susceptible to intranasal infection with VACV (Schoggins et al., 2014) and to footpad inoculation with ectromelia virus, a mouse-specific poxvirus (Wong et al., 2019). Vaccinia B2R gene was recently discovered to encode a cytosolic cGAMP nuclease (renamed as poxin), and deletion of B2R from VACV resulted in attenuation in a skin scarification model (Eaglesham et al., 2019). However, whether poxviruses encode a direct inhibitor(s) of cGAS remains unknown.

In this study, we performed a screen of 82 vaccinia viral genes for inhibition of the cGAS/STING pathway using a dual-luciferase reporter assay and identified several vaccinia genes encoding proteins involved in down-regulating the cGAS/STING/IFNB pathway. Here we show that E5 (encoded by the E5R gene), a BEN-domain-containing protein conserved among orthopoxviruses, is a virulence factor and a major inhibitor of cGAS. E5 interacts with cytoplasmic cGAS and triggers its degradation in a proteasome-dependent manner.

## RESULTS

### Screening strategy for identifying viral inhibitors of the cGAS/STING pathway

Vaccinia virus is a large cytoplasmic DNA virus with a 190 kilobase pairs (kbp) genome that encodes over 200 proteins. MVA has an approximately 30-kbp deletion from its parental vaccinia genome, resulting in the loss of many immune-modulatory viral genes (Antoine et al., 1998). MVA infection of bone marrow-derived dendritic cells (BMDCs) induced IFN-β secretion and cGAMP production, whereas wild-type vaccinia (WT VACV) infection failed to do so (Figures 1A and 1B). These results suggest that WT VACV might encode a viral inhibitor(s) to block cGAS activation and downstream IFN-β production.

**Figure 1.**
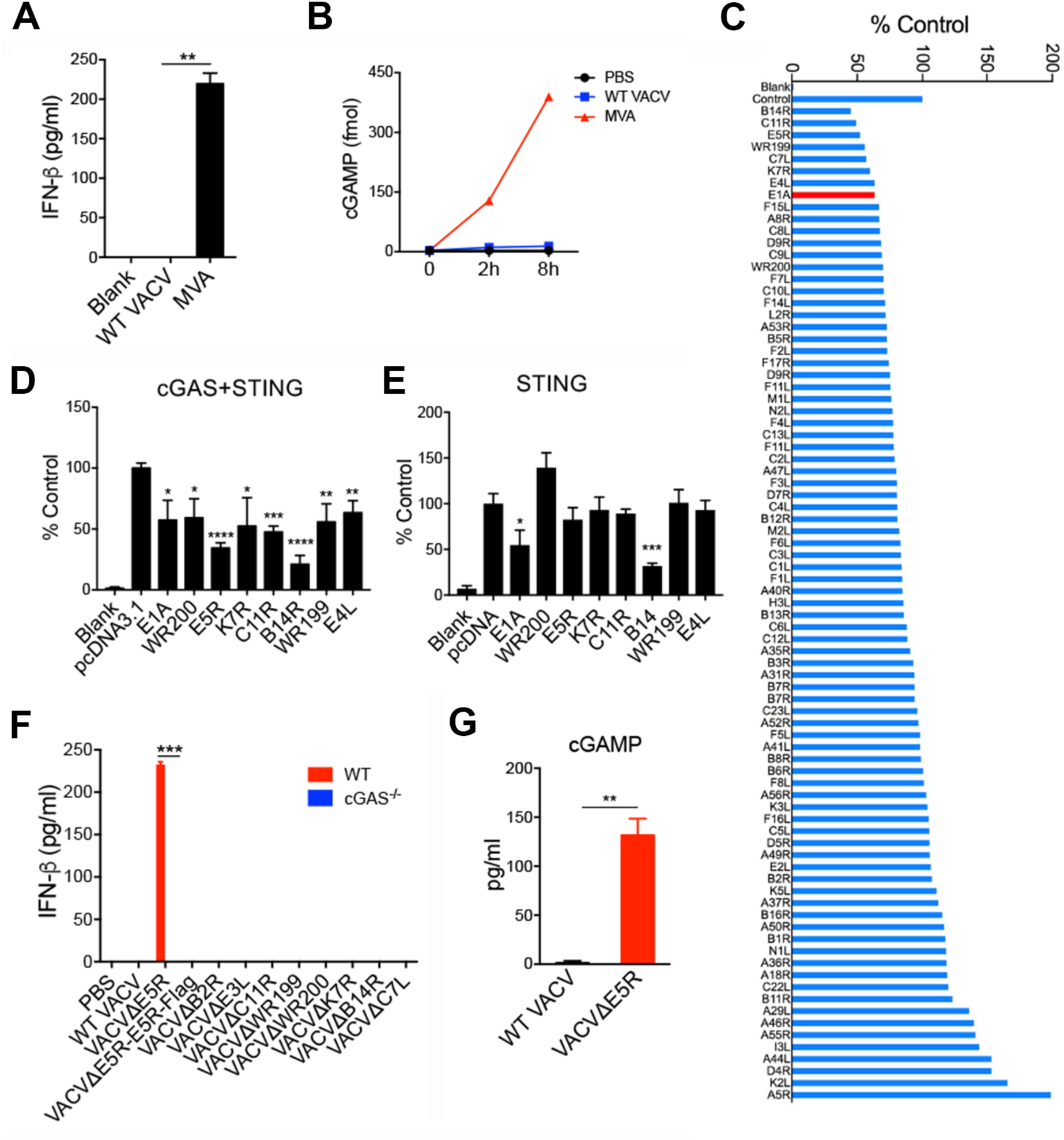
Discovery of vaccinia virus E5 as a key inhibitor of the cGAS-dependent type I IFN pathway. (A) ELISA analysis of IFN-β levels in the supernatants of BMDCs infected with either WT VACV or MVA at a MOI of 10 for 16 h. (B) cGAMP levels in BMDCs infected with either WT VACV or MVA at a MOI of 10. Cells were harvested at 2 and 6 h post infection. cGAMP levels were measured by LC-MS. (C) A dual-luciferase assay to screen for vaccinia viral inhibitors of the cGAS-STING-mediated type I IFN pathway. HEK293T cells were transfected with an IFNB-firefly luciferase reporter, a control plasmid pRL-TK expressing *Renilla* luciferase, cGAS and STING-expressing plasmids, individual vaccinia protein-expressing plasmid, or pcDNA3.1 control plasmid. Adenovirus E1A-expressing plasmid was used as a positive control. Cells were harvested at 24 h post-transfection, and luminescence was determined and expressed as % control. (D) Same as C. A dual-luciferase assay to verify potential vaccinia viral inhibitors of the cGAS-STING pathway. (E) A dual-luciferase assay to verify potential vaccinia viral inhibitors of the STING-IFNB pathway. Similar to D, except that the cGAS-expressing plasmid was not co-transfected with STING-expressing plasmid. (F) ELISA analysis of IFN-β levels in BMDCs from WT or *cGas^-/-^* mice infected with different vaccinia viruses at MOI of 10 for 16 h. (G) ELISA analysis of cGAMP levels in BMDCs from WT mice infected with either WT VACV or VACV*Δ*E5R at MOI of 10 for 16 h. *\** p<0.05, *\*\** p<0.01, *\*\*\** p<0.001 and *\*\*\*\** p<0.0001 (unpaired t test).

To identify potential cGAS inhibitors from the vaccinia genome, we first selected 82 viral genes, mainly the ones expressed at early times during vaccinia infection (Yang et al., 2010), with the reasoning that antagonists of innate immunity are likely encoded by viral early genes. A dual-luciferase assay was then performed to screen for the abilities of these viral proteins to inhibit the cGAS/STING-mediated cytosolic DNA-sensing pathway. Briefly, HEK293T-cells were transfected with plasmids expressing an IFNB-firefly luciferase reporter, a pRL-TK control plasmid expressing *Renilla* luciferase, murine cGAS, human STING, and individual vaccinia viral genes as indicated (Figure 1C). Adenovirus E1A, which inhibits this pathway via interaction with STING (Lau et al., 2015), was used as a positive control for this screening assay (Figure 1C). This assay identified several vaccinia viral early genes (E5R, K7R, B14R, C11R, WR199/B18R, WR200/B19R, E4L) as potential inhibitors of the cGAS/STING pathway (Figures 1C and 1D). Over-expression of all of the candidates except B14R had little effects on STING-induced IFNB promoter activity, suggesting that B14 might target STING or its downstream signaling pathways while other candidates might target cGAS (Figure 1E). Among these genes, WR200/B19R is known to encode a type I IFN binding protein (Symons et al., 1995). Although K7, B14, and C11 have been described as vaccinia virulence factors (Benfield et al., 2013; Chen et al., 2008; Martin et al., 2012), and WR199/B18R was reported to encode a host range factor (Liu et al., 2018; Sperling et al., 2009), how they evade the type I IFN pathway is unclear. E4L encodes vaccinia RNA polymerase subunit RPO30 and a intermediate transcription factor (Ahn et al., 1990; Rosales et al., 1994), and is essential for vaccinia life cycle. Whereas E5 was reported to be a viral early protein associated with the virosomes (viral factories) (Murcia-Nicolas et al., 1999), whether or not it plays a role in immune evasion is unknown.

### Deleting the E5R gene from the WT VACV genome results in cGAS-dependent type I IFN induction in DCs

We hypothesized that deleting a major inhibitor of the cGAS/STING pathway from the VACV genome would result in higher induction of type I IFN than other deletion mutants or the parental virus. To test this idea, we generated a series of recombinant VACV viruses with deletions of individual candidate viral inhibitors, including VACVΔE5R, VACVΔB2R, VACVΔE3L, VACVΔC11R, VACVΔWR199, VACVΔWR200, VACVΔK7R, VACVΔB14R, and VACVΔC7L. Among them, only VACVΔE5R could induce IFN-β secretion from WT BMDCs, but not from cGAS^-/-^ BMDCs (Figure 1F). VACVΔE5R-E5R-Flag in which E5R was replaced by E5R-Flag failed to induce IFN-β secretion, signifying that E5R-Flag is biologically active (Figure 1F). Moreover, whereas B2 (encoded by the B2R gene) was identified as a cGAMP nuclease (Eaglesham et al., 2019), we found that VACVΔB2R infection did not induce IFN-β secretion from WT BMDCs (Figure 1F), suggesting the presence of other viral inhibitors of the cGAS/STING/IFNB pathway in VACVΔB2R. In addition, VACVΔE5R infection of WT BMDCs induced cGAMP production, indicating cGAS activation (Figure 1G). These results demonstrate that vaccinia E5R encodes a major inhibitor of cGAS.

### The vaccinia E5R gene, which encodes a BEN-domain protein, is conserved among orthopoxviruses

Vaccinia E5 is a 341-amino acid polypeptide, comprising an N-terminal alpha-helical domain (amino acids 60-106) and two BEN domains at the C-terminus (amino acid 112-222 and amino acid 233-328) (Figure S1A). BEN was named for its presence in BANP/SMAR1, poxvirus E5R, and NAC1 (Abhiman et al., 2008), and BEN domain-containing proteins function in DNA binding, chromatin organization, and transcriptional repression (Dai et al., 2013; Fedotova et al., 2019; Sathyan et al., 2011). E5R is conserved among orthopoxviruses (Figure S1B and S1C) but less so among yatapoxviruses and myxoma virus. However, E5R orthologs are absent in parapoxviruses, entomopoxviruses, fowlpox, molluscum contagiosum virus (data not shown). The E5 proteins from vaccinia (Western Reserve and Copenhagen), variola (the causative agent for smallpox), and cowpox viruses contain an extra 10-amino acid sequence at the N-termini compared with E5 proteins from vaccinia (Ankara), MVA, and ectromelia (mousepox) (Figure S1C). Interestingly, Monkeypox E5 has large deletions at both its N- and C-termini (Figure S1C, (Douglas and Dumbell, 1996).

### Vaccinia virus E5 is a virulence factor

To test whether vaccinia E5 is a virulence factor, we performed an intranasal infection experiment with WT VACV or VACVΔE5R (2 x 10^6^ pfu) in WT C57BL/6J mice. All of the mice infected with WT VACV lost weight quickly, starting on the third day of infection, and either died or were euthanized due to more than 30% weight loss at days 7 to 8 post-infection (Figures 2A and 2B). By contrast, mice infected with VACVΔE5R lost close to 15% of initial body weight on average at day 6 post-infection, and then recovered (Figures 2A and 2B). These results indicate that VACVΔE5R is attenuated compared with WT VACV and thereby demonstrate that E5 is a virulence factor. Furthermore, VACVΔE5R (2 x 10^7^ pfu) gained virulence in *cGas*^-/-^, or *Sting^gt/gt^* mice but remained attenuated in *Mda5^-/-^* mice, indicating the cytosolic DNA-sensing pathway mediated by cGAS or STING is indispensable for host defense against intranasal infection with VACVΔE5R (Figure 2C and 2D). Moreover, we detected IFN-β in the bronchoalveolar lavage (BALF) 48 h after VACVΔE5R infection (Figure 2E), indicating that intranasal infection with VACVΔE5R infection could induce IFN-β production in vivo.

**Figure 2.**
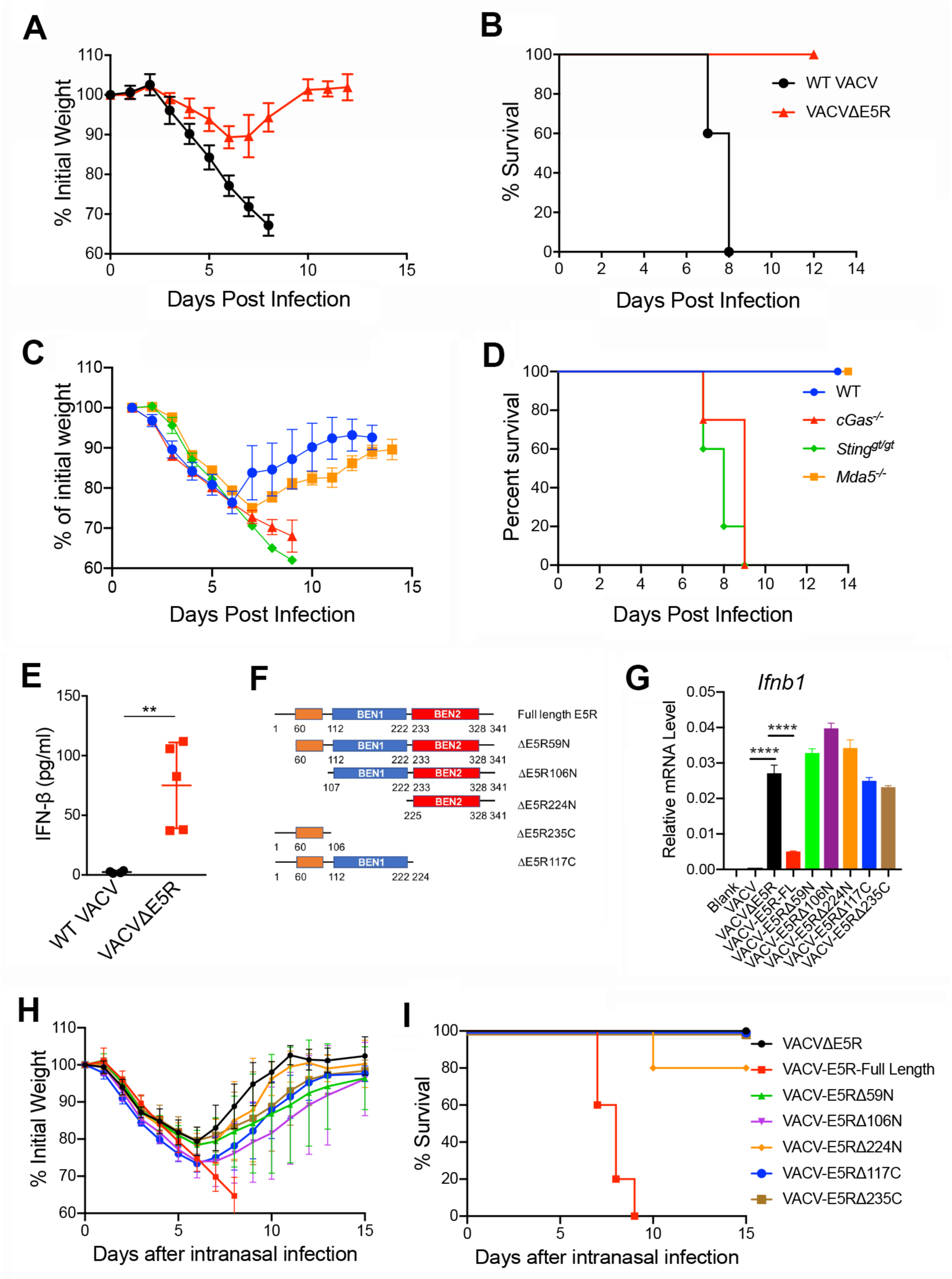
Vaccinia virus E5 is a virulence factor in vivo. (A-B) Percentages of initial weight (A) and Kaplan-Meier survival curves (B) of WT C57BL/6J mice (n=5 in each group) over days post-intranasal infection with either WT VACV or VACVΔE5R at a dose of 2 x 10^6^ pfu per mouse. (C-D) Percentages of initial weight (C) and Kaplan-Meier survival curves (D) of WT, *Mda5^-/-^*, *cGas^-/-^*, or *Sting^gt/gt^* mice (n=5 in each group) over days post-intranasal infection with VACVΔE5R at a dose of 2 x 10^7^ pfu per mouse. (E) ELISA analysis of IFN-β levels in the BALF harvested from WT mice at 48 h post-infection with WT VACV or VACVΔE5R at a dose of 2 x 10^7^ pfu per mouse. (F) Schematic diagram of VACV E5R full-length revertant and various VACV E5R truncation mutants. (G) RT-PCR analysis of *Ifnb* levels in WT BMDCs infected with different vaccinia viruses including VACV, VACVΔE5R, VACV-E5R-FL revertant, and various VACV E5R truncation mutants for 6 h. (H-I) Percentages of initial weight (H) and Kaplan-Meier survival curves (I) of WT C57BL/6J mice (n=5 in each group) over days post-intranasal infection with VACVΔE5R, VACV-E5R-full length revertant, and various E5R truncation mutants a dose of 2 x 10^7^ pfu per mouse. *\*\** p<0.01 and *\*\*\*\** p<0.0001 (unpaired t test).

To determine which domain(s) of E5 are required for E5-mediated inhibition of IFNB induction and virulence, we constructed VACV-E5R (full-length) and a series of its truncation mutants, as shown in Figure 2F. Whereas VACV-E5R (full-length) only mildly induced IFNB gene expression in BMDC cells (Figure 2G), all E5R truncation mutants induced IFNB gene expression at a similar level to VACVΔE5R (Figure 2G). Moreover, intranasal infection of VACV-E5R (full-length) and E5R truncation mutants (2 x 10^7^ pfu) in C57BL/6J mice showed that only VACV-E5R (full-length) infection was lethal, while all of the truncation mutants caused transient weight loss but 100% survival, except for VACV-E5RΔ224N (with 80% survival) (Figure 2H and 2I). These results demonstrated that all E5 domains, including the N-terminal alpha-helical domain and the two BEN domains, are required for repression of Ifnb gene expression and virulence.

### Deleting E5R from the MVA genome strongly induces cGAMP production and type I IFN secretion in BMDCs

To investigate whether the E5R gene of the MVA genome encodes a functional protein, we generated MVAΔE5R. MVAΔE5R infection of BMDCs potently upregulated *Ifnb1*, *Ifna*, *Ccl4*, and *Ccl5* gene expression (Figure 3A), whereas MVA infection had modest induction. MVAΔE5R infection of BMDCs induced much higher levels of IFN-β secretion than MVA, or heat-inactivated MVA (heat-iMVA), or heat-inactivated MVAΔE5R (heat- iMVAΔE5R) (Figure 3B). In addition, heat-iMVAΔE5R caused higher levels of IFN-β secretion than heat-iMVA (Figure 3B). Heat-inactivation at 55°C for one hour prevents viral protein expression in infected cells (Dai et al., 2017). Therefore the difference in IFN-β induction between heat-iMVAΔE5R and heat-iMVA-infected DCs might be attributed to virion E5 protein brought into the cells by heat-iMVA. IFN-β induction in BMDCs depended on viral doses (Figure S2A), and that IFN-*α* secretion was also strongly induced by MVAΔE5R infection in BMDCs (Figure 3C). Consistent with that, MVAΔE5R caused much higher levels of cGAMP production in BMDCs than MVA (Figure 3D). Similar to BMDC, MVAΔE5R infection of bone marrow-derived macrophages (BMM) or plasmacytoid DCs (pDCs) also induced much higher levels of IFN-β secretion than MVA (Figure S2B and S2C). These results demonstrated that vaccinia E5 blocks cGAS activation, and deletion of E5R from the MVA genome potently activates the cGAS/STING pathway in multiple myeloid cell types. Furthermore, MVAΔE5R-induced IFN-β secretion from BMDCs was abolished in cGAS^-/-^ or STING^Gt/Gt^ cells and diminished in IRF3^-/-^ or IRF7^-/-^ cells (Figure 3E). In addition to BMDCs, MVAΔE5R-induced IFN-β secretion in BMMs, pDCs, or primary fibroblasts was also dependent on the cGAS/STING pathway (Figures S2D-S2F). These results demonstrate that the cGAS/STING-mediated cytosolic DNA-sensing pathway and the transcription factors IRF3 and IRF7 are required for MVAΔE5R-induced IFN-β secretion in various primary cell types.

**Figure 3.**
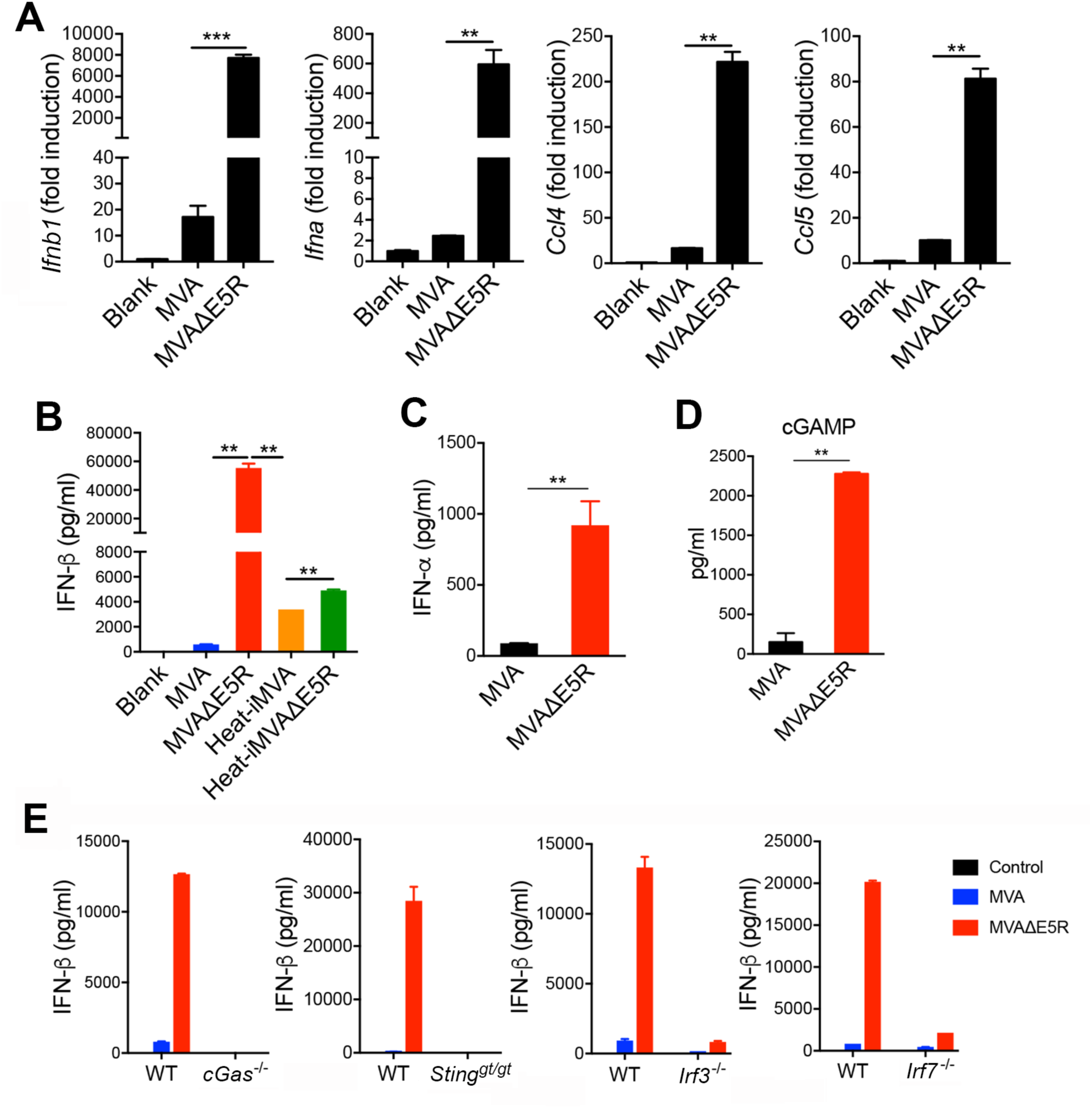
MVAΔE5R strongly induces type I IFN production in a cGAS/STING, IRF3/IRF7-dependent manner. (A) RT-PCR of *Ifnb1*, *Ifna*, *Ccl4*, and *Ccl5* gene expression in WT BMDCs infected with either MVA or MVAΔE5R at a MOI of 10 for 6 h. (B-C) ELISA analyses of IFN-β or IFN-*α* levels in the supernatants of WT BMDCs infected with MVA, MVAΔE5R, Heat-iMVA or Heat-iMVAΔE5R at a MOI of 10 for 16 h. (D) ELISA analyses of cGAMP levels in WT BMDCs infected with MVA or MVAΔE5R at a MOI of 10 for 16 h. (E) ELISA analyses of IFN-β levels in the BMDC from WT, *cGas^-/-^*, *Sting^gt/gt^*, *Irf3^-/-^*, and *Irf7^-/-^* mice infected with MVA or MVAΔE5R at a MOI of 10 for 16 h. *\*\** p<0.01, *\*\*\** p<0.001 and *\*\*\** p<0.001 (unpaired t test).

In addition to parental viral DNA provided by the incoming virions, progeny viral DNA generated after DNA replication in the virosomes may also stimulate the cytosolic DNA sensor cGAS, resulting in IFN-β production. To evaluate this, we used phosphonoacetate (PAA) or aphidicolin to block viral DNA replication (DeFilippes, 1984; Moss and Cooper, 1982). PAA or aphidicolin treatment of MVAΔE5R-infected MEFs blocked virosome formation as expected (Figure S2G), and abolished the expressions of viral late genes such as A27 and A34 (Figure S2H). Both *Ifnb1* and *Ifna* gene expressions induced by MVAΔE5R were partially reduced in the presence of PAA or aphidicolin (Figure S2I). Overall, our results indicate both the parental and progeny viral DNA from MVAΔE5R-infected BMDCs contribute to type I IFN induction.

### WT VACV or MVA infection triggers cGAS degradation via a proteasome-dependent mechanism

To investigate how E5 antagonizes the cGAS/STING pathway, we first evaluated cGAS protein levels after WT VACV infection. We observed that cGAS protein levels were lower at six hours after WT VACV infection in BMDCs compared with mock-infection control (Figure 4A), suggesting that cGAS protein might be degraded after viral infection. Treatment with proteasome inhibitor MG132 prevented cGAS degradation, whereas treatment with a pan-caspase inhibitor, Z-VAD, or an AKT1/2 inhibitor VIII had little effect on cGAS levels (Figure 4A). Treatment with the protein translation inhibitor cycloheximide (CHX) partially blocked cGAS degradation, suggesting that the newly synthesized viral proteins might facilitate cGAS degradation (Figure 4A). These results indicate that WT VACV-induced cGAS degradation is proteasome-dependent. Unlike WT VACV, VACVΔE5R infection of BMDCs did not result in cGAS degradation (Figure 4B). Similarly, whereas MVA infection of BMDCs triggered cGAS degradation, MVAΔE5R infection did not (Figure 4C), confirming that E5 contributes to cGAS degradation in the context of either VACV or MVA infection. Similar to what we observed with VACV, MVA-induced decline of cGAS levels was fully reversed by MG132 and partially reversed by CHX (Figure 4D). In contrast, PAA did not affect MVA-caused reduction of cGAS levels, suggesting that the degradation of cGAS is independent of viral DNA replication (Figure 4D).

**Figure 4.**
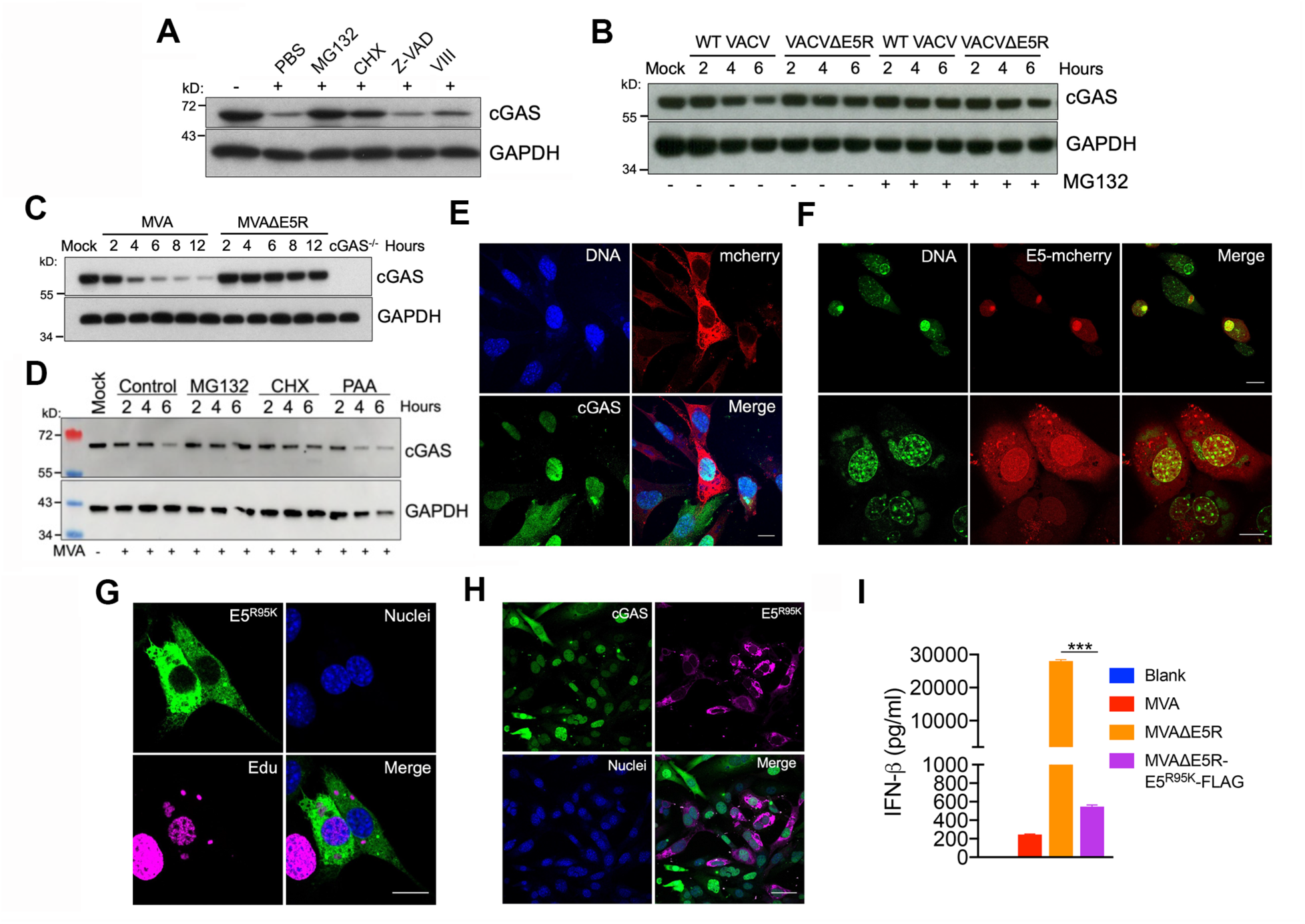
WT VACV or MVA infection induces proteasome-dependent degradation of cGAS. (A) Immunoblot of cGAS in MEFs infected with WT VACV for 6 h. Cells were pretreated with either cycloheximide (CHX, 25 µg/ml), a proteasome inhibitor MG132 (25 µM), a pan-caspase inhibitor Z-VAD (50 µM), an AKT1/2 inhibitor VIII (10 µM) for 30 min and then infected with WT VACV in the presence of each individual drug. Cells were collected at 6 h post-infection. (B) Immunoblot of cGAS in BMDCs infected with either WT VACV or VACV*Δ*E5R. Cells were pre-treated with or without MG132 (25 µM) for 30 min and infected with WT VACV or VACVΔE5R in the presence or absence of MG132. Cells were collected at 2, 4 and 6 h post-infection. (C) Immunoblot of cGAS in BMDCs infected with MVA or MVA*Δ*E5R. (D) Immunoblot of cGAS in BMDC infected with MVA. Cells were pre-treated with MG132 (25 µM), CHX (25 µg/ml), PAA (200 µg/ml) or PBS for 30 min and then infected with MVA at a MOI of 10 in the presence of each drug or PBS control. Cells were collected at 2, 4 and 6 h post-infection. (E) Representative confocal images showing GFP-cGAS protein in GFP-cGAS MEFs cells after MVA-mCherry infection for 6 h. Scale bar, 15 μm. (F) Representative confocal images showing E5-mCherry expression in BMDCs (top) or MEFs (bottom) after MVA-E5R-mCherry infection at a MOI 10 for 6 h. DNA staining with SiR-DNA dye highlights nuclei and virosomes. Scale bar, 15 μm. (G) Representative confocal images showing E5^R95K^-Flag expression in MEFs after MVA*Δ*E5R-E5^R95K^-Flag infection for 6 h. Nuclear and virosomal DNAs were stained EdU. Scale bar, 15 μm. (H) Representative confocal images showing E5^R95K^-Flag and cGAS in GFP-cGAS MEFs after MVA*Δ*E5R-E5^R95K^-Flag infection for 6 h. Scale bar, 15 μm. (I) ELISA analyses of IFN-β levels in supernatants of BMDCs infected with MVA, MVAΔE5R or MVA*Δ*E5R-E5^R95K^-Flag at a MOI of 10 for 16 h.

Using MEFs that express GFP-cGAS, in which GFP was tagged to the N-terminus of cGAS, we observed that at 6 h after MVA-mCherry infection, cytoplasmic cGAS was not detectable in mCherry^+^ MVA-infected cells. However, nuclear cGAS remained in those infected cells, suggesting that MVA infection triggered the degradation of only cytoplasmic cGAS (Figure 4E). By contrast, in mCherry^-^ uninfected cells, cGAS was degraded neither in the cytoplasm nor in the nucleus (Figure 4E). To better understand vaccinia E5 localization, we constructed a recombinant MVA expressing vaccinia E5-mcherry (MVA-E5R-mCherry), in which the C-terminus of vaccinia E5 is fused with mCherry for live-cell imaging. At 6 h post-infection with MVA-E5-mCherry, E5 was detected in both cytoplasms and nuclei of infected BMDCs (Figure 4F, upper panel) and MEFs (Figure 4F, lower panel). In the process of generating an MVAΔE5R-E5R-Flag virus, we isolated one mutant strain, MVAΔE5R-E5^R95K^-Flag, which contains a single nucleotide change (G284A), resulting in the replacement of arginine at amino acid 95 of E5 by lysine. E5^R95K^-Flag was expressed in the cytoplasms but was not in the nuclei of MVAΔE5R-E5^R95K^-Flag-infected MEFs (Figure 4G). Infection of MEFs expressing GFP-cGAS with MVAΔE5R-E5^R95K^-Flag resulted in the degradation of cGAS in the cytoplasm but not in the nucleus (Figure 4H). In addition, IFN-β production was diminished in BMDCs infected with MVAΔE5R-E5^R95K^-Flag (Figure 4I), indicating that the cytoplasmic E5 is sufficient to induce cGAS degradation and to suppress type I IFN production.

### Vaccinia virus E5 protein interacts with cGAS and promotes K48-linked poly-ubiquitination of cGAS and subsequent degradation

To test whether E5 alone can trigger cGAS degradation without viral infection, we co-transfected a cGAS-expressing plasmid with an E5R-expressing plasmid or empty vector. 24 h later, cells were infected with MVA*Δ*E5R at a MOI of 10 for 6 h in the presence or absence of CHX. We observed that the cGAS level was decreased after co-transfection with the former but not the latter (Figure 5A and S3A). Moreover, MVAΔE5R infection failed to enhance E5-mediated cGAS degradation in the presence or absence of CHX (Figure 5A), indicating that E5 alone can trigger cGAS degradation, likely in the context of plasmid transfection.

**Figure 5.**
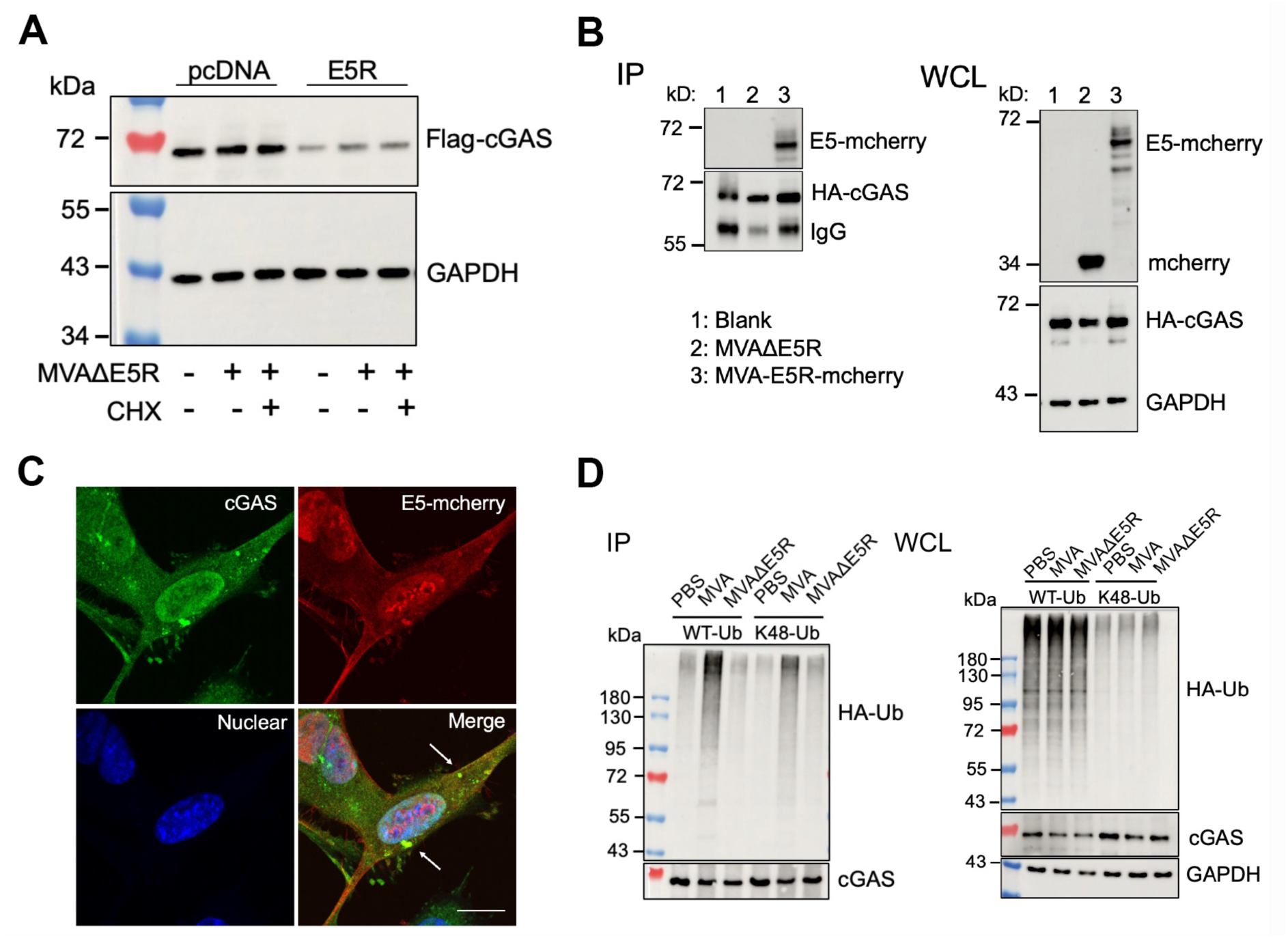
E5 interacts with cGAS and promotes K48-linked poly-ubiquitination of cGAS. (A) HEK293T cells were co-transfected with Flag-cGAS and pcDNA-E5R or pcDNA3.1 plasmids. 24 h later, cells were infected with MVA*Δ*E5R at a MOI of 10 for 6 h in the presence or absence of CHX (25 µg/ml). Flag-cGAS protein levels were determined by an anti-Flag antibody. (B) HEK293T cells were transfected with an HA-cGAS-expressing plasmid. 24 h later, cells were infected with either MVA*Δ*E5R or MVA*Δ*E5R-E5R-mCherry at a MOI of 10 for 6 h in the presence of MG132 (25 µM). HA-cGAS was pulled down by an anti-HA antibody and E5-mCherry was determined by an anti-mCherry antibody. (C) Representative confocal images showing cGAS and E5-mCherry co-localization in the cytoplasm of MEF-cGAS-GFP cells after MVA-E5R-mCherry infection at MOI 10 for 6 h in the presence of MG132 (25 µM). Scale bar, 15 μm. White arrows point to some representative puncta of E5 and cGAS co-localization in the cytoplasm. (D) HEK293T cells were co-transfected with V5-cGAS and HA-Ub (WT or K48 only)- expressing plasmids. 24 h later, cells were infected with MVA or MVA*Δ*E5R at MOI 10 for 6 h at the presence of MG132 (25 µM). cGAS was pulled down by an anti-V5 antibody and ubiquitinated cGAS was determined by an anti-HA antibody. IP: immunoprecipitation. WCL: whole cell lysates.

To assess whether E5 and cGAS interact with each other, we transfected HEK293T cells with HA-cGAS expression plasmid and then infected with MVAΔE5R (in which mcherry was expressed independently of E5) or MVA-E5R-mCherry (in which mCherry was tagged to the E5 C-terminus) in the presence of MG132. Immunoprecipitation with anti-HA antibody pulled down E5-mCherry but not mCherry, thus indicating an E5-cGAS interaction (Figure 5B). Confocal imaging of MEFs expressing cGAS-GFP infected with MVA-E5R-mCherry virus in the presence of MG132 showed that E5 and cGAS co-localization to punctate cytoplasmic structures (Figure 5C).

We next hypothesized that vaccinia E5 induces cGAS ubiquitination and subsequent proteasome-dependent degradation. We detected higher levels of cGAS ubiquitination, particularly K48-linked poly-ubiquitination, in cells infected with MVA compared with those infected with MVAΔE5R (Figure 5D). Thus, our results support that E5 expressed by MVA promotes K48-linked poly-ubiquitination of cGAS, leading to its degradation.

To test whether E5 binds to DNA, we used an *in vitro* transcription/translation system including reticulocyte lysate, T7 RNA polymerase, and ^35^S-methionine. Radio-labeled cGAS or E5 proteins were tested for DNA binding by using DNA-coupled beads. Both cGAS and E5 could be pulled down individually with DNA-coupled beads but not by beads alone without DNA (Figures S3B). When cGAS and E5 were expressed together, both proteins could be pulled down by DNA-coupled beads. These results suggest E5 is capable of binding DNA. However, we cannot rule out whether cGAS and E5 compete for DNA binding in this assay, because DNA beads were used in excess (Figures S3B).

### E5 is ubiquitinated in MVA-infected cells

To evaluate whether E5 is ubiquitinated in MVA-infected cells, we first used Halo-4xUBA^UBQLN1^ beads, which contain four tandem ubiquitin-associated (UBA) domains from Ubiquilin-1 to specifically pull down ubiquitinated proteins (Ordureau et al., 2014) in MVA-infected cells (Figure 6A). Here we show that E5 was among the top viral proteins with a high % of coverage and high peptide numbers (Figure 6B). Other viral proteins that are enriched for ubiquitination include RNA polymerase subunits (A24, J6), viral DNA replication factors (I3, D5, E9), ribonucleotide reductase subunits (I4 and F4), and some immunomodulatory proteins (E5, B15, C16, E3, and C7). We next engineered a recombinant MVA with a human immunoglobulin (IgG) Fc domain-tagged to the C-terminus of E5, showing that E5-Fc protein was pulled down by protein A agarose. Subsequently, ubiquitinated E5 was determined by western blot using an anti-HA antibody, thus suggesting that E5 is ubiquitinated after viral infection (Figure 6C).

**Figure 6.**
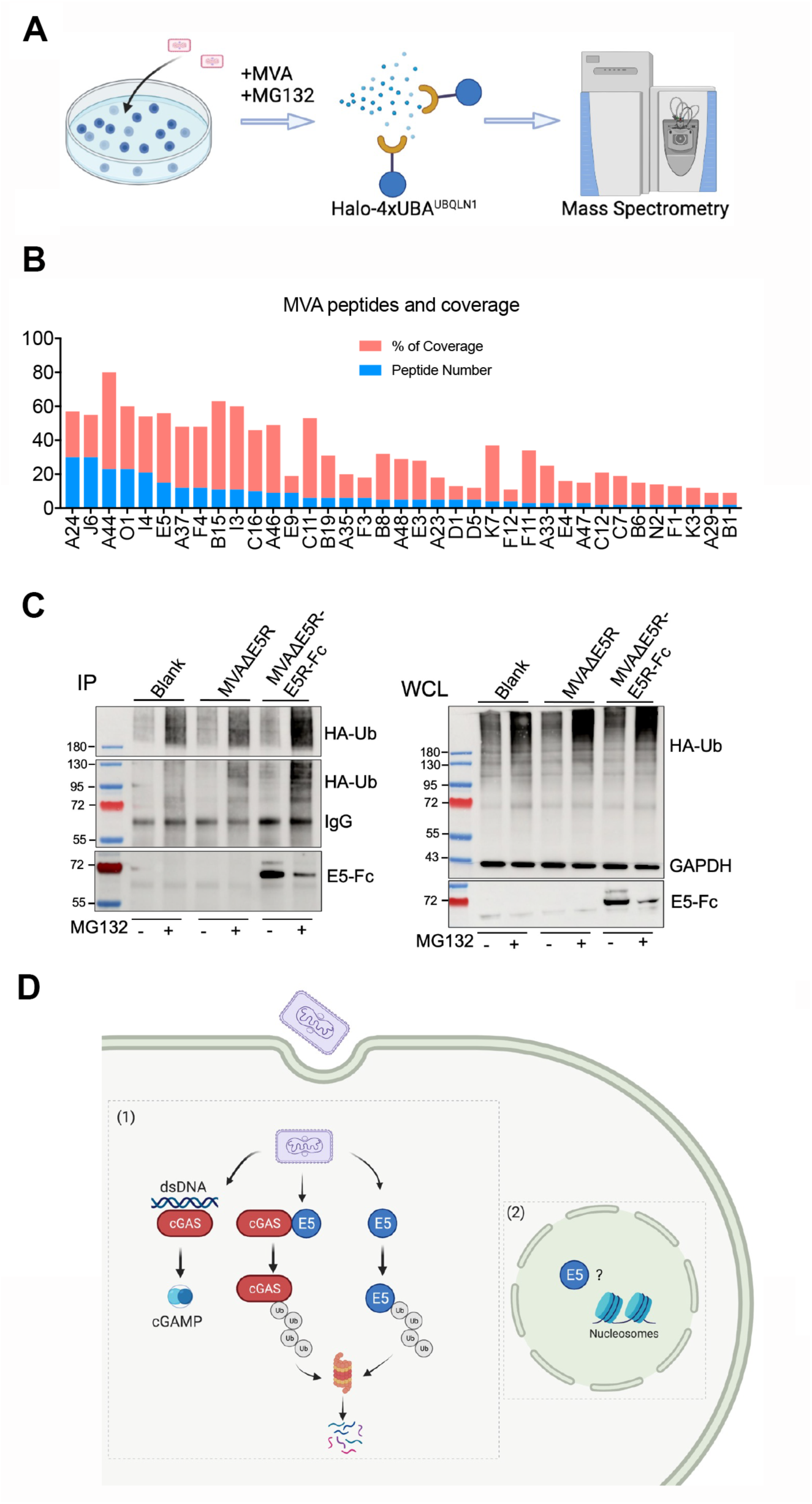
E5 protein is ubiquitinated after MVA infection and working model. (A) Schematic diagram of the experimental design. In brief, HEK293T cells were infected with MVA at a MOI 10 for 6 h in the presence of MG132 (25 µM). Ubiquitinated proteins were pulled down by Halo-4xUBA^UBQLN1^ beads, which have high affinity to ubiquitinated proteins. After further purification, peptides bound to beads were measured by mass spectrometry. (B) Peptide numbers and coverage of MVA peptides from mass spectrometry. (C) HEK293T cells were transfected with HA-Ub (WT)-expressing plasmid. 24 h later, cells were infected with MVA*Δ*E5R or MVA*Δ*E5R-E5-Fc at a MOI of 10 for 6 h. Cells were treated with or without MG132 (25 µM) during infection. E5-Fc was pulled down by protein A agarose and ubiquitinated E5 was determined by an anti-HA antibody.

### Deleting the E5R gene from MVA improves the immunogenicity of the vaccine vector

MVA has been investigated as a vaccine vector for various infectious diseases and cancers, and MVA infection modestly activates human monocyte-derived DCs (moDCs) (Drillien et al., 2004).

To investigate whether E5R deletion improves the immunogenicity of the viral vector, we first generated MVAΔE5R-OVA, expressing a model antigen chicken ovalbumin (OVA) and then compared DC maturation upon MVAΔE5R-OVA vs. MVA-OVA infection. We observed that MVAΔE5R-OVA infection induced higher levels of CD86 and CD40 expression compared with MVA-OVA at 24 h post-infection (Figure 7A-C). However, both MVAΔE5R-OVA and MVA-OVA-induced CD86 and CD40 expression diminished in cGAS^-/-^ BMDCs, indicating that the cytosolic DNA-sensing pathway is essential for MVAΔE5R-OVA or MVA-OVA-induced DC maturation (Figure 7A-C).

**Figure 7.**
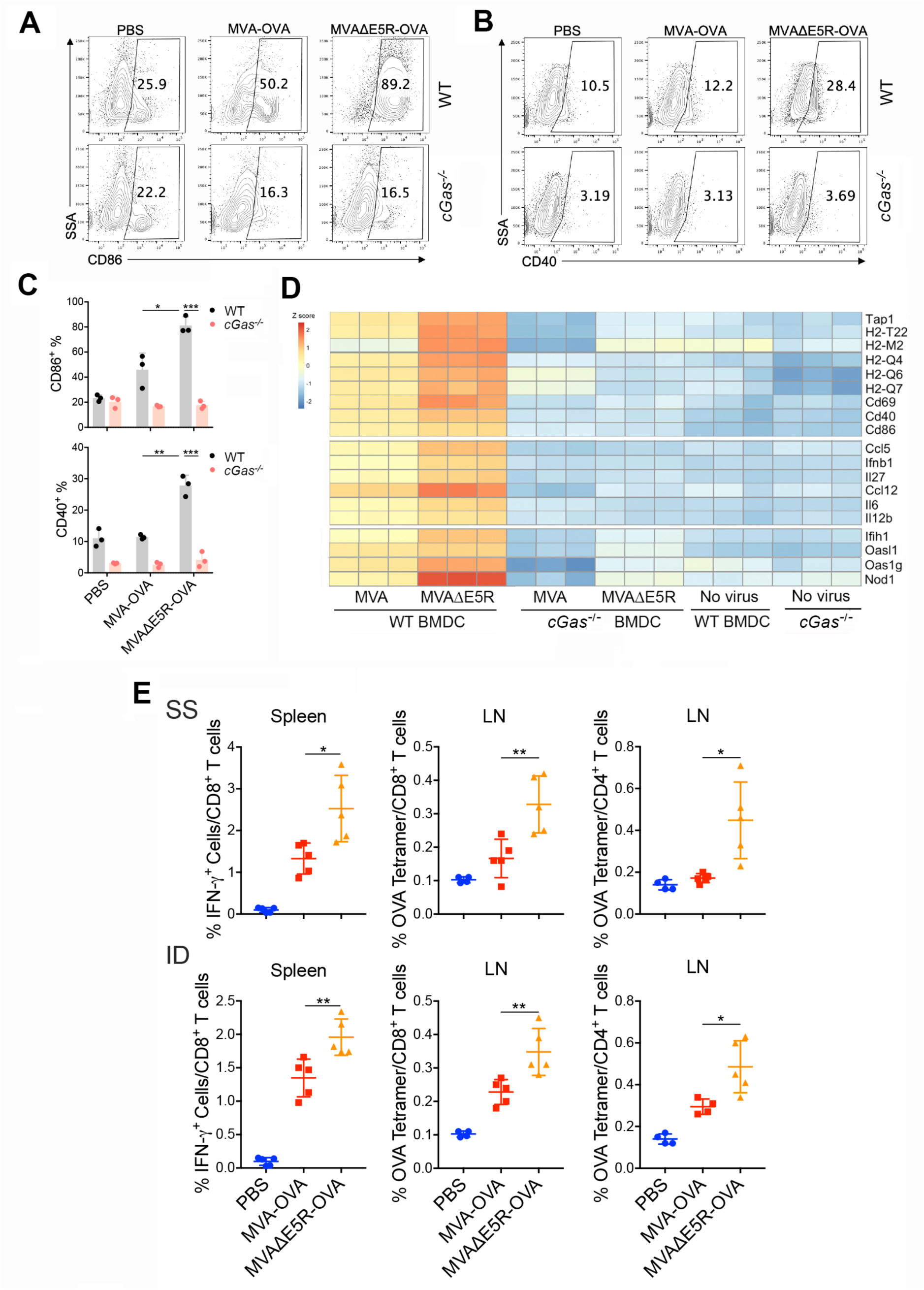
MVAΔE5R-OVA promotes DC maturation and antigen-specific CD8^+^ T cells activation. (A-C) Representative flow cytometry dot plots (A, B) or analysis (C) of CD86 and CD40 expression in WT or *cGas*^-/-^ BMDC infected with MVA-OVA or MVA*Δ*E5R-OVA at MOI 10 for 16 h. (D) Heatmap showing relative expression of selected immune-related genes in WT or *cGas*^-/-^ BMDC infection with MVA or MVAΔE5R. These include genes involved in antigen presentation, DC activation, IFN and proinflammatory cytokines and chemokines, and innate immune sensors. (E) Antigen-specific T cell responses after vaccination with MVA-OVA or MVAΔE5R-OVA. C57BL/6J mice were vaccinated with MVA-OVA or MVA*Δ*E5R-OVA via skin scarification (SS) or intradermal injection (ID). One week later, spleens and draining lymph nodes (dLNs) were harvested and SIINFEKL-specific CD8^+^ T cells in splenocytes and OVA tetramer-specific CD8^+^ or CD4^+^ T cells in lymph nodes were determined by FACS. *\** p<0.05, *\*\** p<0.01 and *\*\*\** p<0.001 (unpaired t test).

RNA-seq analysis of WT or cGAS^-/-^ BMDCs infected or mock-infected with either MVA or MVAΔE5R demonstrated that MVAΔE5R infection in WT BMDCs induced higher levels of type I IFN and pro-inflammatory cytokines and chemokines genes, including *Ifnb*, *Ccl5*, *Ccl12*, *Il12b*, *Il6*, *Il27*, DC maturation and activation markers such as *CD86*, *CD40*, and *CD69*, as well as genes involved in antigen cross-presentation, including *Tap1*, *H2-Q4*, *H2-Q6*, and *H2-Q7*, compared with MVA (Figure 7D, S4A and S4B). The upregulation of these genes by both MVA and MVAΔE5R was cGAS-dependent (Figure 7D and S4C).

Next, we performed vaccination through skin scarification (SS) or intradermal (ID) injection with either MVA-OVA or MVAΔE5R-OVA. One week after vaccination, anti-OVA CD8^+^ and CD4^+^ T cells in the spleens and draining lymph nodes were analyzed. Vaccination with MVAΔE5R- OVA resulted in more OVA-specific CD8^+^ T cells in the spleens than MVA-OVA (Figure 7E). And more OVA-specific CD8^+^ or CD4^+^ T cells were detected in the draining lymph nodes (dLN) after MVAΔE5R-OVA vaccination, compared with MVA-OVA(Figure 7E).

## DISCUSSION

The identification of vaccinia E5 as a major inhibitor of the cytosolic DNA-sensor cGAS highlights the importance of that pathway in host defense against poxvirus infection. E5, a founding member of the BEN-domain family, is conserved among orthopoxviruses. Here we show that the virulence factor E5 binds to cGAS, triggering cGAS ubiquitination and proteasome-dependent degradation, and that deleting E5R from the MVA viral vector improves its immunogenicity.

Virulent poxviruses, including VACV (Western Reserve and Copenhagen strains), cowpox, and ectromelia virus, fail to activate STING, unlike the highly attenuated derivative, MVA (Dai et al., 2014; Georgana et al., 2018). In addition, MVA but not WT VACV infection of BMDCs induces cGAMP production, suggesting that VACV encodes an inhibitor(s) of cGAS. Through an unbiased screen of 82 vaccinia genes, we identified several candidate genes that might encode cGAS inhibitors, including E5R, K7R, C11R, WR199/B18R, and WR200/B19R. We then focused on E5R, because a VACV mutant lacking E5R induced IFN-β secretion and cGAMP production in BMDCs, while VACV mutants lacking other individual candidate genes failed to induce IFN-β secretion in BMDCs. In our study, VACV lacking the B2R gene, which encodes a cGAMP nuclease, fails to induce type I IFN production in BMDC, suggesting that there might be an additional vaccinia viral protein(s) antagonizing the cGAS/STING pathway.

Although E5 was first identified by mass spectrometry as one of the three major early viral proteins associated with virosomes in vaccinia-infected cells (Murcia-Nicolas et al., 1999), the function of E5 remained elusive. E5 was also found in the highly purified virions by mass spectrometry after chemical crosslinking, and several interaction partners were identified, including RNA polymerase subunits RAP94, RP147, and NTP1 (Mirzakhanyan and Gershon, 2019). Here we show E5 presence in both the nuclei and cytoplasm of the infected cells. In MVAΔE5R-E5^R95K^-Flag-infected cells, E5^R95K^ localizes only to the cytoplasm, but is sufficient for mediating cGAS degradation and IFN inhibition. The R95K mutation is within a putative nuclear localization signal of E5, ^92^KFKRMIR^98^.

We show that E5 mediates cGAS degradation via a proteasome-dependent pathway. We propose the following working model based on our results (Figure 6D). Vaccinia virus enters host cells via micropinocytosis (Mercer and Helenius, 2008). Upon viral entry, viral DNA is detected by the cytosolic DNA sensor cGAS, whose activation leads to cGAMP production and subsequent STING stimulation. However, in the presence of vaccinia E5, which is mainly synthesized by the incoming virions as an early viral protein, cGAS is targeted for ubiquitination and proteasome-dependent degradation through interacting with E5. This leads to reduced cGAMP production and *Ifnb1* gene expression. Some of the newly expressed E5 is ubiquitinated and degraded in the cytoplasm, while some of the E5 pool localizes to the nucleus. Although the function of nuclear cGAS is inhibited by nucleosomes (Boyer et al., 2020; Kujirai et al., 2020), E5 does not target nuclear cGAS for degradation. The function of nuclear E5 needs further investigation.

We previously reported that viral replication is not important for MVA sensing in BMDCs (Dai et al., 2014). In this study, however, MVAΔE5R-induced Ifnb gene expression was partially reduced in the presence of the viral DNA replication inhibitors, PAA and aphidicolin. This result suggests that virosomal progeny viral DNA is detected by cGAS in the setting of MVAΔE5R infection.

Various post-translational modifications of cGAS have been reported, including ubiquitination, phosphorylation, acetylation, sumoylation, glutamylation, neddylation, and caspase-mediated cleavage (Song et al., 2020; Wu and Li, 2020). For example, RNF185, a RING domain E3 ubiquitin ligase, has been shown to interact with cGAS during human simplex virus-1 (HSV-1) infection, stimulating K27-linked poly-ubiquitination of cGAS, important for cGAS enzymatic activity (Wang et al., 2017). TRIM56, an IFN-inducible E3 ubiquitin ligase, interacts with cGAS to promote monoubiquitination and cGAS activity (Seo et al., 2018). In addition, TRIM14, an IFN inducible protein, recruits USP14 to cleave K48-linked poly-ubiquitin chains of cGAS and thereby inhibiting cGAS degradation (Chen et al., 2016). The ubiquitin E3 ligase responsible for K48- linked poly-ubiquitination remains elusive. Here we show that MVA infection induces K48-linked poly-ubiquitination of cGAS and promotes its degradation in a proteasome-dependent manner. E5 is critical for this process via interacting with cGAS. However, the exact details of how E5 recruits a viral or cellular E3 ubiquitin ligase to catalyze K48-linked poly-ubiquitination of cGAS remains to be determined.

Despite these limitations, the discovery of E5 as a major inhibitor of cGAS provides significant insights into improving MVA as a vaccine vector. Here we show that MVAΔE5R-OVA infection of BMDCs induces high levels of IFN-β production and DC maturation, consistent with cGAS-dependent transcriptomic changes induced by MVAΔE5R. Vaccination with MVAΔE5R-OVA induces higher levels of OVA-specific CD8^+^ and CD4^+^ T cells compared with MVA-OVA. Recent studies have shown that MVA-based vaccine vectors expressing SARS-CoV-2 spike protein induce potent anti-spike T and B cell immune responses and provides protection in animal models (Garcia-Arriaza et al., 2021; Liu et al., 2021; Routhu et al., 2021; Tscherne et al., 2021). Future investigations of whether MVAΔE5R-based vaccine vectors improve vaccine efficacy against infectious agents such as SARS-CoV-2 is warranted.

## Acknowledgements

We thank the Flow Cytometry Core Facility and Molecular Cytology Core Facility at the Sloan Kettering Institute. We thank Drs. Charles Rice and Stewart Shuman for helpful discussions. We thank Curt Balch for editing. This work was supported by NIH grants K-08 AI073736 (L.D.), R56AI095692 (L.D.), R03 AR068118 (L.D.), and MSK Technology Development Fund (L.D.), sponsored research agreement from IMVAQ Therapeutics (L.D.). Cancer Prevention and Research Institute of Texas, RP180725 (Z.J.C.). Z.J.C. is an investigator of the Howard Hughes Medical Institute. This research was also funded in part through the NIH/NCI Cancer Center Support Grant P30 CA008748.

## Author Contributions

Author contributions: N.Y. and L.D. were involved in all aspect of this study, including conceiving the project, designing and performing experiments, data analyses and interpretation, manuscript writing. N.Y. designed the screen for vaccinia inhibitors of the cytosolic DNA-sensing pathway and performed most of the experiments. Y.W. assisted N.Y. in the screen and validation of vaccinia inhibitors. Y.W and L.D. performed RNA-seq experiments. Y.W. and L.D. generated various VACV E5 deletion viruses and performed pathogenesis studies. P.D. and T.L. are involved in cGAMP measurement in BMDCs infected by vaccinia and MVA. A.T, T.Z., and J.Z.X. analyzed RNA-seq data, and assisted in manuscript preparation. C.Z. and H.F. are involved the cGAS and E5 DNA-binding assay. H.P., Z.L, R.H., and A.O. are involved LS/MS determination of ubiquination viral proteins. A.O., C.Z., H.F., and Z.J.C. assisted in experimental design, data interpretation, and manuscript preparation. L.D. provided overall surpervision of the project.

## MATERIALS AND METHODS

### Mice

Female C57BL/6J mice between 6 and 8 weeks of age were purchased from the Jackson Laboratory and were used for the preparation of bone marrow-derived dendritic cells (BMDCs) and for intranasal infection experiments. *cGas^-/-^* mice were purchased from the Jackson Laboratory. *Sting^gt/gt^* mice were generated in the laboratory of Russell Vance (University of California, Berkeley) (Sauer et al., 2011). *Mda5^-/-^* mice were generated in Marco Colonna’s laboratory (Washington University) (Gitlin et al., 2006). These mice were maintained in the animal facility at the Sloan Kettering Cancer Institute. All procedures were performed in strict accordance with the recommendations in the Guide for the Care and Use of Laboratory Animals of the National Institute of Health and the protocol was approved by the Committee on the Ethics of Animal Experiments of Sloan-Kettering Cancer Institute.

### Viruses

The Western Reserve (WR) strain of vaccinia virus (VACV) was propagated, and virus titers were determined on BSC40 (African green monkey kidney cells) monolayers at 37°C. MVA and MVA-OVA viruses were kindly provided by Gerd Sutter (University of Munich) and propagated in BHK-21 (baby hamster kidney cell, ATCC CCL-10) cells. All viruses were purified through a 36% sucrose cushion. Heat-iMVA or Heat-iMVAΔE5R was generated by incubating purified MVA or MVAΔE5R virus at 55 °C for 1 hour. To generate recombinant VACVΔE5R virus, BSC40 cells were infected with WT vaccinia virus (WR) at a MOI of 0.2. After 1-2 h, cells were transfected with pE5R-mCherry plasmids with Lipofectamine 2000 (Invitrogen). Homologous recombination between the plasmid DNA and vaccinia viral genome resulted in the deletion of *E5R* gene from the viral genome and insertion of mCherry, under the control of the vaccinia synthetic early and late promoter (pSE/L). Cells were collected two days later and underwent three cycles of freeze-thaw. Plaque purification was performed based on the mCherry fluorescence seen under the microscope. After 4-5 rounds, pure recombinant VACVΔE5R-mCherry viruses were obtained, and validation of E5R deletion confirmed by PCR analyses and DNA sequencing. Various other deletion mutants, including VACVΔB2R, VACVΔE3L, VACVΔC11R, VACVΔWR199, VACVΔWR200, VACVΔK7R, VACVΔB14R, and VACVΔC7L were generated following a procedure similar to that described above. VACVΔE5R-E5R-Flag was generated by inserting the E5R-Flag sequence into the *TK* locus of VACVΔE5R. VACV-E5R-FL, VACV-E5RΔ59N, VACV-E5RΔ106N, VACV-E5RΔ224N, VACV-E5RΔ117C and VACV-E5RΔ235C were generated by inserting full-length or truncated E5R into the *E5R* locus of VACVΔE5R. MVAΔE5R and MVAΔE5R-OVA expressing mCherry was generated through homologous recombination at the E4L and E6R loci flanking *E5R* gene of the MVA or MVA-OVA genome in BHK21 cells following a procedure similar to that described above. MVAΔE5R-E5R-Fc was generated by inserting E5R-Fc sequence into the *TK* locus of MVAΔE5R using fluorescence color selection. MVA-E5R-mCherry was generated by inserting E5R-mCherry sequence into the *TK* locus of MVA. MVAΔE5R-E5^R95K^-Flag was generated by inserting E5R-Flag sequence into the *TK* locus of MVAΔE5R using drug selection. The recombinant virus was enriched in the presence of gpt selection medium including MPA, xanthine and hypoxanthine, and plaque purified for at least four rounds. During the selection process, a spontaneous mutation occurred resulting in the generation of MVAΔE5R-E5^R95K^-Flag, verified by PCR and DNA sequencing.

### Intranasal infection of vaccinia virus in mice

Female C57BL/6J mice between 6 and 8 weeks of age (5-10 in each group) were anesthetized and infected intranasally with WT VACV, VACVΔE5R or VACVΔE5 expressing E5R full-length revertant and various E5R truncation mutants at the indicated doses in 20 µl PBS. Mice were monitored and weighed daily. Those that lost over 30% of their initial weight were euthanized. Kaplan-Meier survival curves were determined. Bronchoalveolar lavage fluid (BALF) was harvested following intratracheal infusion of 1 ml of cold PBS.

### Vaccination with MVA-OVA or MVAΔE5R-OVA

6 to 8-week-old Female C57BL/6J mice were vaccinated via skin scarification (SS) or intradermal (ID) injection with MVA-OVA or MVAΔE5R-OVA at a dose of 2x10^7^ pfu. One week later, spleens and draining lymph nodes (dLNs) were collected and processed using the Miltenyi GentleMACS™ Dissociator. Splenocytes were stimulated with OVA_257-264_ (SIINFEKL) peptide (5 μg/ml). After 1 h of stimulation, GolgiPlug (BD Biosciences) (1:1000 dilution) was added and incubated for 12 h. Cells were then treated with BD Cytofix/Cytoperm™ kit prior to staining with respective antibodies for flow cytometry analyses. The antibodies used for this assay are as follows: BioLegend: CD3e (145-2C11), CD4 (GK1.5), CD8 (53-5.8), IFN-*γ* (XMG1.2).

dLNs were digested with collagenase D (2.5 mg/ml) and DNase (50 µg/ml) at 37°C for 25 min before filtering through 70-µm cell strainer. For tetramer staining, cells were incubated with tetramers for 30 mins at 37 °C. Alexa Fluor 647 H-2K(b) ova 257-264 SIINFEKL tetramer and PE I-A(b) Ova 329-337 AAHAEINEA tetramer were synthesized from NIH Tetramer Core Facility. Cells were analyzed on the BD LSRFortessa.

### Cell lines and primary Cells

BSC40, HEK293T, MEFs, and cGAS-GFP MEFs (Yang et al., 2017) were cultured in Dulbecco’s modified Eagle’s medium supplemented with 10% fetal bovine serum (FBS), 2 mM L-glutamine and 1% penicillin-streptomycin. BHK-21 were cultured in Eagle’s Minimal Essential Medium (Eagle’s MEM, Life Technologies, Cat# 11095-080) containing 10% FBS, and 1% penicillin-streptomycin. For the generation of GM-CSF-BMDCs, bone marrow cells (5 million cells in each 15 cm cell culture dish) were cultured in RPMI-1640 medium supplemented with 10% fetal bovine serum (FBS) in the presence of GM-CSF (20 ng/ml, BioLegend, Cat# 576304) for 9-12 days as described in (Dai et al., 2014). For the generation of bone marrow-derived macrophages (BMMs), bone marrow cells were cultured in RPMI-1640 medium supplemented with 10% fetal bovine serum (FBS) in the presence of M-CSF (10 ng/ml, PeproTech, Cat# 315-02) for 7-9 days.

To culture primary murine plasmacytoid dendritic cells (pDCs), bone marrow cells were cultured in RPMI-1640 medium supplemented with 10% FBS in the presence of FMS-like tyrosine kinase 3 ligand (Flt3L) (100 ng/ml, R&D Systems, Cat# 308-FK) for 7-9 days. Cells were fed every 2–3 days by replenishing 50% of the medium. pDCs were gated as B220^+^PDCA-1^+^ cells and sorted in a FACS Aria II instrument (BD Biosciences) as described in (Dai et al., 2011). The antibodies used for this assay are as follows: B220 (RA3-6B2) and PDCA-1 (927) from Biolegend.

Mice epidermal sheet were removed as previously described (Deng et al., 2008). Briefly, skins were washed with cold Ca^2+^ and Mg^2+^ free PBS and then incubated in a digestion buffer containing 1 U dispase/ml at 37°C for 1 h. Epidermal sheets were mechanically removed, the remaining dermis was washed in Ca^2+^ and Mg^2+^ free PBS 5 times and incubated in a digestion buffer containing 2 mg/ml collagenase A (Roche), 100 µg/ml of DNase I (Sigma; d4527) and 1% BSA in Ca^2+^ and Mg^2+^ free PBS at 37°C for 1-2 hours. The resulting suspension was filtered through a 100-, 70- and 40-mm nylon mesh sequentially (VWR) and washed two times with a buffer (Ca^2+^ and Mg^2+^ free PBS containing 1% BSA and 2 mM EDTA). Cells were cultured RPMI-1640 medium supplemented with 10% fetal bovine serum (FBS). Only the adherent cells were used after 2-3 days of culture.

### Cytokine assays

The IFN-β levels in BALF were determined using mouse IFN beta ProQuantum Immunoassay kit (ThermoFisher). IFN-*α* and IFN-β levels in the supernatants of cultured BMDCs were determined by ELISA (PBL Biomedical Laboratories).

### Flow cytometry analysis for DC maturation

GM-CSF-BMDCs from WT or cGAS^-/-^ mice were infected with either MVA-OVA or MVAΔE5R-OVA at a MOI of 10 for 16 h. Cells were washed with MACS buffer (Miltenyi Biotec) and stained with antibodies against CD40 (3/23, Biolegend) and CD86 (GL-1, Biolegend). FACS analyses were performed using LSRFortessa^TM^ Cell Analyzer (BD Biosciences). Data were analyzed with FlowJo software (version 10.5.3).

### Immunofluorescence imaging

Cultured cells were plated in Lab-Tek^TM^ II chamber slide (ThermoFisher) and fixed in 4% (w/v) paraformaldehyde at room temperature (RT) for 10 min, permeabilized with 0.5% (v/v) Triton X-100 in PBS for 5 min, and blocked in 5% goat serum (Sigma), 3% bovine serum albumin (Fisher), and 0.1% Triton X-100 at room temperature for 1 hr. Primary antibodies were incubated at 4 °C at the indicated dilutions overnight: chicken anti-GFP (1:1000, Abcam), rat anti-mCherry (1:1000, ThermoFisher), and mouse anti-Flag (1:1000, Sigma). After three washes in PBS, slides were incubated with indicated secondary antibodies, including goat anti-chicken Alexa Fluor-488 (1:1000, Invitrogen), goat anti-mouse Alexa Fluor-488 (1:1000, Invitrogen), or goat anti-rat Alexa Fluor-594 (1:1000, Invitrogen), at RT for 60 min. After three washes in PBS, slides were mounted in ProLong Gold Antifade Mountant (ThermoFisher). Following incorporation of 5-ethynyl-2′-deoxyuridine (EdU) (PMID: 18272492), cells were fixed, permeabilized, and labeled with AF647 azide, according to the Click-iT EdU Imaging Kit protocol (ThermoFisher). Images were acquired using a confocal microscope (Leica TCS SP8 or Zeiss LSM880).

### Live cell imaging

For time-lapse imaging of E5-mCherry expression in MEFs, cells were seeded on Lab-Tek^TM^ II chamber slide (ThermoFisher), and infected with MVA-E5R-mCherry at MOI 10 and stained with fluorogenic SiR-DNA (Cytoskeleton). Cells were incubated at 37°C supplemented with 5% CO_2_. Images were acquired using a ZEISS Axio Observer Z1. All the images were further processed with Image J software.

### Plasmid Construction

IFN-β reporter plasmid (pIFN-β-luc) (Wies et al., 2013) was provided by Michaela Gack (University of Chicago). pRL-TK was purchased from Promega. Human STING expression plasmid was provided by Tom Maniatis (University of Columbia). Murine STING (mSTING) sequences were amplified by PCR and were cloned into pcDNA3.2-DEST plasmids. Human cGAS (hcGAS) and murine cGAS (mcGAS) plasmids were purchased from Invivogen. Flag-cGAS and V5-cGAS were amplified by PCR and were cloned into pcDNA3.2-DEST plasmids. pRK-HA-Ubiquitin-WT, pRK-HA-Ubiquitin-K48, and pBabe 12S E1A were purchased from Addgene. 82 selected VACV genes were amplified by PCR from the VACV WR genome and subcloned into pcDNA3.2-DEST using the Gateway cloning method (Invitrogen).

### The Dual-Luciferase reporter assay

The firefly and *Renilla* luciferase activities were measured using the Dual-Luciferase Reporter Assay system according to the manufacturer’s instructions (Promega). To screen for vaccinia inhibitors of the cGAS/STING pathway, mcGAS (50 ng) and hSTING (10 ng) expression plasmids, together with pIFN-β-luc (50 ng), pRL-TK (10 ng), as well as selected vaccinia gene expression plasmid or adenovirus E1A expression plasmid (200 ng) were transfected into HEK293T cells. 24 h post-transfection, cells were collected and lysed. To assess the effects of vaccinia inhibitors of the STING pathway, mSTING (50 ng) expression plasmid, together with pIFN-β-luc (50 ng), pRL-TK (10 ng), as well as selected vaccinia gene expression constructs or adenovirus E1A expression plasmid (200 ng) were transfected into HEK293T cells. The relative luciferase activity was expressed as arbitrary units by normalizing firefly luciferase activity under the IFNB promoter to *Renilla* luciferase activity from a control plasmid, pRL-TK.

### Western blot analysis

Cells were lysed in RIPA lysis buffer supplemented with 1x Halt™ Protease and Phosphatase Inhibitor Cocktail (ThermoFisher). Protein samples were separated by SDS-PAGE and then transferred to nitrocellulose membrane and incubated with primary antibodies specific for cGAS (CST, 31659), His (ThermoFisher, MA1-21315), FLAG (Sigma, F3165), HA (ThermoFisher, 71-5500), HA (Sigma, H3663), mCherry (ThermoFisher, M11217), and GAPDH (CST, 2118) were used. HRP-conjugated anti-rabbit, mouse, or rat IgG antibody were used as secondary antibodies (CST, 7074, 7076 or 7077). Detection was performed using SuperSignal^TM^ Substrates (Thermo Fisher, 34577 or 34095).

### Co-immunoprecipitation

For cGAS ubiquitination assays, HEK293T cells in 10-cm plates were transfected with V5-cGAS together with HA-Ub-WT or HA-Ub-K48. 24 h later, cells were infected with MVA or MVAΔE5R at MOI 10 for 6 h in the presence of MG132 (25 μg/ml) and lysed in RIPA lysis buffer (ThermoFisher, 89901) on ice for 30 min. Anti-V5 antibody (ThermoFisher, R960-25) was added into cell lysate to a final concentration of 1 μg/ml and incubated at 4 °C overnight on a rotator. The next day, protein G-magnetic beads (Bio-Rad, 161-4023) were added and incubated at 4 °C for 2 h. The beads were washed five times with RIPA buffer. Lastly, the bead-bound proteins were denatured in SDS buffer by heating at 98 °C for 5 min before loading on an SDS-PAGE gel. E5 ubiquitination assays were performed following a procedure similar to those described above. Generally, after HEK293T cells were transfected with HA-Ub, they were then infected with MVAΔE5R or MVAΔE5R-E5R-Fc at MOI 10 for 6 h. After cell lysis with RIPA buffer, protein A-Agarose beads (ThermoFisher, 20333) were added and incubated at 4 °C for 2 h, washed five times, and the bead-bound proteins were denatured in SDS buffer by heating at 98 °C for 5 min before loading on an SDS-PAGE gel.

### Quantitative real-time PCR

Total RNA was extracted from whole cell lysates using TRIzol reagent (Invitrogen) or with RNeasy Plus Mini kit (Qiagen). RNAs were reverse-transcribed and amplified by PCR using the Verso cDNA synthesis kit (Thermo Fisher) and SYBR^TM^ Green Master Mix (Thermo Fisher). Cellular RNAs were normalized to GAPDH levels. All assays were performed on an ABI 7500 system and analyzed with ABI 7500 SDS software v.1.3 (Applied Biosystems). Primer sequences used are listed in Table S1.

### DNA-coupled beads binding assay

Beads coupled to single-end biotinylated DNA were generated as previously described (Postow et al., 2008). DNA-coupled beads and uncoupled beads were washed twice in binding buffer (10 mM Tris-Cl pH 7; 80 mM NaCl; 0.05 % Triton X-100; 1 mM DTT). ^35^S methionine-labeled full-length cGAS and E5 was generated with the TnT Coupled Reticulocyte Lysate System (Promega) according to the manufacturer’s instructions. TnT reactions without added template DNA served as control. TnT reactions were diluted 1:10 in binding buffer supplemented with BSA to a final concentration of 0.04 μg/μl, and beads coupled to 1 µg of DNA were incubated in 20 μl of this mixture at 20 °C for 45 min under agitation. Beads were then washed three times in binding buffer, and bound proteins were eluted with SDS sample buffer, and analyzed by gel electrophoresis, followed by scanning on a phosphoimager of the dried gel.

### cGAMP measurement by liquid chromatography-mass spectrometry (LC-MS)

cGAMP was measured by LC-MS as previously reported (Li et al., 2021). Briefly, cell pellets were supplemented with 80 fmol internal standard (^15^N_10_-cGAMP, in-house generated) and were extracted subsequently in 80% methanol and 2% acetic acid, and twice in 2% acetic acid to obtain metabolite extract. cGAMP was enriched from combined extracts on HyperSep Aminopropyl SPE Columns (Thermo Scientific). After washing twice in 2% acetic acid and once in 80% methanol, samples were eluted in 4% ammonium hydroxide in 80% methanol. Vacuum-dried eluents were dissolved in water and analyzed on a Dionex U3000 HPLC coupled with TSQ Quantiva Triple Quandruple mass spectrometer (Thermo Scientific). The chromatography used LUNA NH_2_ resin (5 µm, Phenomenex) as stationary phase packed in 0.1 mm ID × 70 mm L silica capillaries. Mobile phases are acetonitrile (A), and 20 mM ammonium bicarbonate and 20 mM ammonium hydroxide aqueous solution (B). Flow rate is 800 nL/min (0-4 min), 300 nL/min (4-19 min), and 600 nL/min (19-27 min), with a gradient of 20% B (0-3 min), 50% B (4 min), 80% B (14-18 min), and 20% B (19-27 min). cGAMP and standard were analyzed by multiple reaction monitoring in the positive mode with the following transitions: 675-136, 675-152, 675-476, and 675-524 for cGAMP; and 685-136, 685-157, 685-480, and 685-529 for the ^15^N_10_-cGAMP standard. Endogenous cGAMP levels were calculated by multiplying the cGAMP-to-standard ratios by 80 fmol (the amount of standard spiked into each sample).

### Detection of ubiquitinated vaccinia viral proteins by LC-MS

To detect ubiquitinated viral protein during vaccinia virus infection, HEK293T cells were infected with MVA at MOI 10 for 6 h in the presence of MG132. Ubiquitinated proteins were purified using Halo-4× UBA^UBQLN1^ as previously described (Ordureau et al., 2014). Briefly, whole-cell extracts (1 mg) that were lysed in lysis buffer containing 100 mM chloroacetamide and incubated at 4 °C for 16 h with 30 μL of Halo-4× UBA^UBQLN1^ beads (pack volume). Following four washes with lysis buffer containing 1 M NaCl, and one final wash in 10 mM Tris (pH 8.0), proteins were released from Halo-4× UBA^UBQLN1^ beads by 6 M guanidine HCL.

For MS, the released proteins were purified by SP3 protocol and digested with 20 µl of trypsin (20 ng/µl) and lysC (10 ng/µl) on beads at 37°C for 2 hours followed by incubation at 24°C overnight. After de-salting, samples were subjected to reduction (10 mM TCEP) and alkylation (20 mM chloroacetamide). After digested overnight at 37 °C with trypsin, and further purified with SP3 protocol (BD Biosciences), samples were analyzed by LC/ MS(Waters nanoAcquity UHPLC/Thermo QExactuve Plus).

The LC-MS/MS .raw files were processed using Mascot (version 2.6.1.100) and searched for protein identification against the SwissProt protein database for human (downloaded January 7^th^, 2020) and vaccinia virus (downloaded September 17^th^, 2021). Carbamidomethylation of C was set as a fixed modification and the following variable modifications allowed: oxidation (M), N-terminal protein acetylation, deamidation (N and Q), ubiquitination (K), and phosphorylation (S, T and Y). Search parameters specified an MS tolerance of 10 ppm, an MS/MS tolerance at 0.080 Da and full trypsin digestion, allowing for up to two missed cleavages. False discovery rate was restricted to 1% in both protein and peptide level. Protein coverage and peptide count were obtained using Scaffold (4.8.4).

### RNA-seq analyses of GM-CSF-cultured BMDCs infected with MVA vs. MVAΔE5R

GM-CSF-cultured BMDCs (1 x 10^6^) from WT or cGAS^-/-^ mice were infected with MVA or MVAΔE5R at a multiplicity of infection (MOI) of 10. Cells were collected at 16 h post-infection. Total RNA was extracted from collected cells using RNeasy Plus Mini kit (Qiagen) according to manufacturer’s protocol. Total RNA integrity was analyzed using a 2100 Bioanalyzer (Agilent Technologies). Messenger RNA was prepared using TruSeq Stranded mRNA Sample Library Preparation kit (Illumina, San Diego, CA) according to the manufacturer’s instructions. The normalized final cDNA libraries were pooled and sequenced on Illumina NovaSeq6000 sequencer with pair-end 50 cycles. The raw sequencing reads in BCL format were processed through bcl2fastq 2.19 (Illumina) for FASTQ conversion and demultiplexing.

The resulting FASTQ files were processed using kallisto (PMID: 27043002) followed by a tximport (PMID: 26925227) transcript-to-gene level transformation. Transcript indices for kallisto were created using FASTA files from Ensembl release 95 for mm10 for all annotated cDNA and ncRNA transcripts, as well as FASTA sequences for WT VACV transcripts (accession no. NC_006998.1). Gene-level read counts were then processed using the limma suite of tools (PMID: **25605792**) first with a voom transformation, followed by linear model fitting to determine differentialy expressed genes, and lastly performing gene set testing using the CAMERA function. Select gene sets were plotted as row normalized z-scores in heatmaps using voom normalized counts.

### Statistics

Two-tailed unpaired Student’s t test was used for comparisons of two groups in the studies. Survival data were analyzed by log-rank (Mantel-Cox) test. The p values deemed significant are indicated in the figures as follows: *, p < 0.05; **, p < 0.01; ***, p < 0.001; ****, p < 0.0001. The numbers of animals included in the study are discussed in each figure legend.

