## Supplemental files for "Vaccinia E5 is a major inhibitor of the DNA sensor cGAS"

This file contains:

- Supplementary Fig. 1
- Supplementary Fig. 2
- Supplementary Fig. 3
- Supplementary Fig. 4
- Supplementary Table 1

#### Figure S1

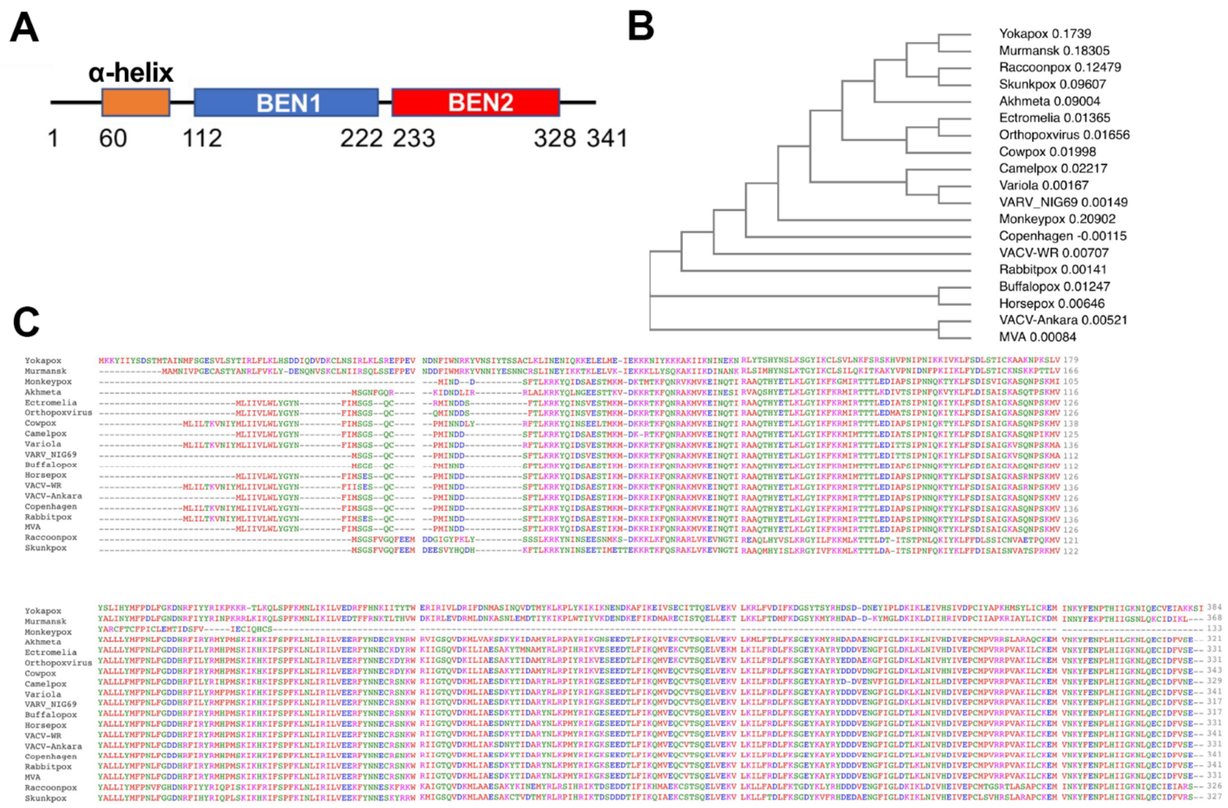


**Figure S1 related to Figure 1. Sequence alignment of E5 from orthopoxviruses.**

(A) Schematic diagram of E5 protein, which is comprised of an N-terminal α-helical and two C-terminal BEN domains.

(B) Phylogenetic tree of E5 sequence alignments.

(C) Multiple sequence alignments of E5 from orthopoxviruses.

### Figure S2

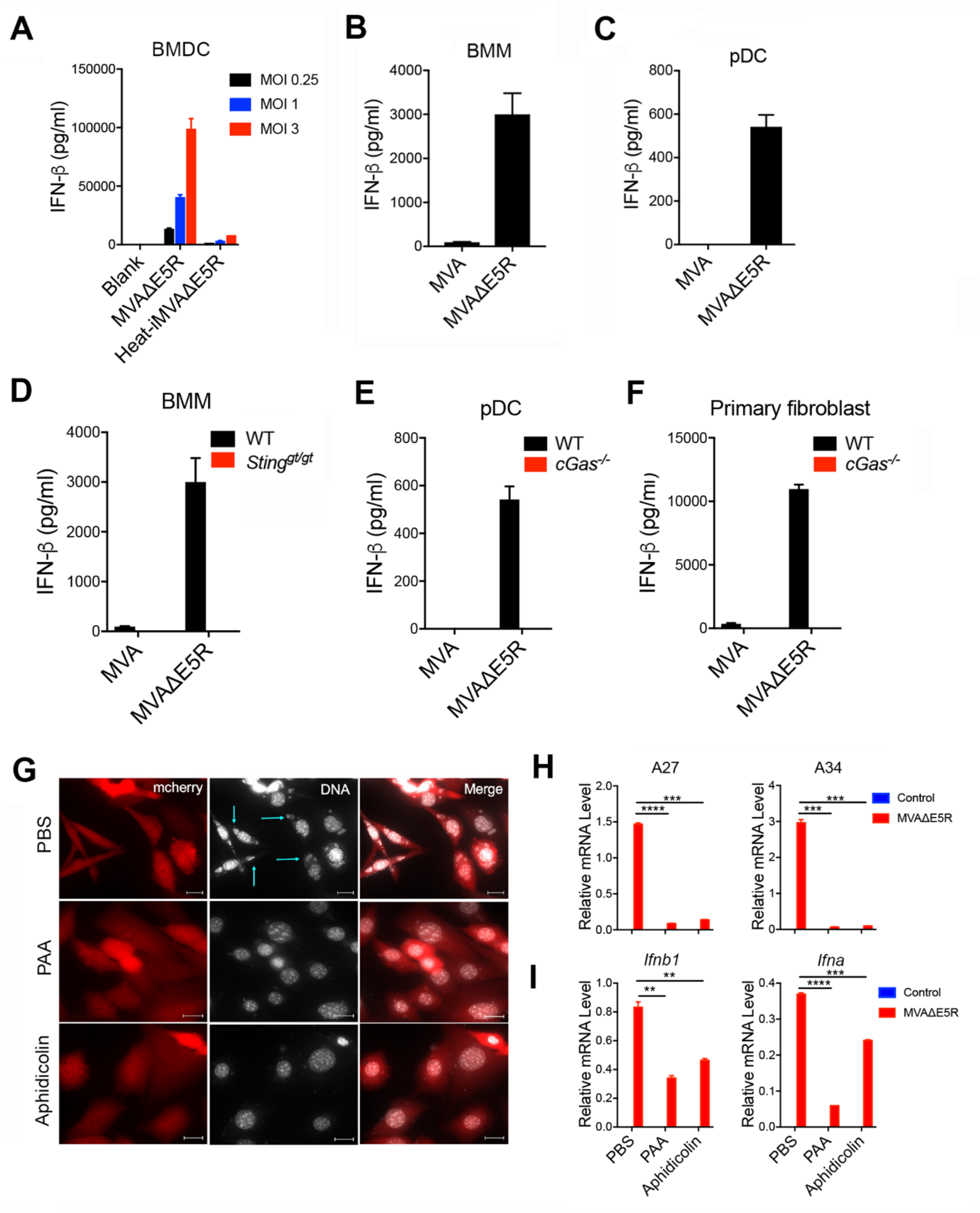
A
BMDC
MOI 0.25
MOI 1
MOI 3
IFN-β (pg/ml)
150000
100000
50000
0
Blank
MVAΔE5R
Heat-iMVAΔE5R
B
BMM
IFN-β (pg/ml)
4000
3000
2000
1000
0
MVA
MVAΔE5R
C
pDC
IFN-β (pg/ml)
800
600
400
200
0
MVA
MVAΔE5R
D
BMM
WT
Sting gt/gt
IFN-β (pg/ml)
4000
3000
2000
1000
0
MVA
MVAΔE5R
E
pDC
WT
cGas-/-
IFN-β (pg/ml)
800
600
400
200
0
MVA
MVAΔE5R
F
Primary fibroblast
WT
cGas-/-
IFN-β (pg/ml)
15000
10000
5000
0
MVA
MVAΔE5R
G
mcherry
DNA
Merge
PBS
PAA
Aphidicolin
H
A27
A34
Control
MVAΔE5R
Relative mRNA Level
2.0
1.5
1.0
0.5
0.0
4
3
2
1
0
****
***
I
Ifnb1
Ifna
Control
MVAΔE5R
Relative mRNA Level
1.0
0.8
0.6
0.4
0.2
0.0
0.4
0.3
0.2
0.1
0.0
**
****
***
PBS
PAA
Aphidicolin

**Figure S2 related to Figure 3. MVAΔE5R strongly induces type I IFN production in a cGAS/STING-dependent manner**

(A) ELISA analyses of IFN-β levels in the supernatants of WT BMDC infected with MVAΔE5R or Heat-iMVAΔE5R at different MOIs for 16 h.

(B-C) ELISA analyses of IFN-β levels in the supernatants of bone marrow-derived macrophages (BMM; A) or plasmacytoid dendritic cells (pDC; B) infected with MVA or MVAΔE5R at a MOI of 10 for 16 h.

(D-F) ELISA analyses of IFN-β levels in the supernatants of WT or $Sting^{gt/gt}$ BMM (D), WT or $cGas^{-/-}$ pDC (E) or WT or $cGas^{-/-}$ primary fibroblasts (F) infected with MVA, or MVAΔE5R at a MOI of 10 for 16 h.

(G) Representative images showing virosome in MEF cells treated with PAA, Aphidicolin or PBS after infection with MVAΔE5R at a MOI of 10 for 6 h. Scale bar, 15 μm.

(H) RT-PCR of *Ifnb1*, *Ifna*, A27, and A34 gene expression of BMDC from WT mice infected with MVAΔE5R or PBS control at a MOI of 10 for 6 h. The cells were treated with PAA, Aphidicolin or PBS during virus infection.

#### Figure S3

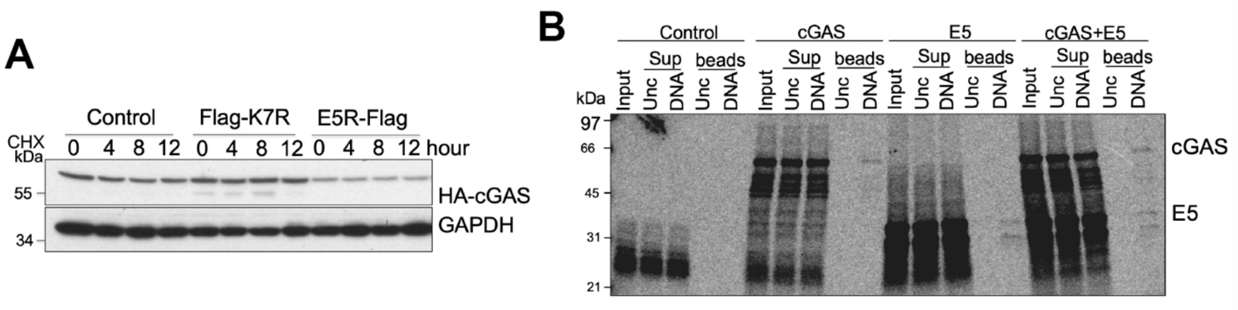


**Figure S3 related to Figure 5. E5 interacts with cGAS and promotes K48-linked poly-ubiquitination of cGAS**

(A) HEK293T cells were co-transfected with HA-cGAS and pcDNA-E5R-flag, pcDNA-Flag-K7R or pcDNA3.1 plasmid. 24 h later, cells were treated with CHX (25 μg/ml) and collect at the indicated time points. HA-cGAS protein level was determined by using an anti-HA antibody.

(B) DNA-beads binding assays using $^{35}$S methionine-labeled-cGAS or E5 from *in vitro* transcription/translation. DNA-beads or uncoupled beads were incubated with $^{35}$S methionine-labeled-cGAS, E5 or cGAS and E5 together. Supernatants (Sup) were separated from the beads using a magnet. The beads were washed extensively, and bound proteins were eluted with reducing SDS sample buffer. Unc: uncoupled beads. DNA: DNA-coupled beads.

### Figure S4

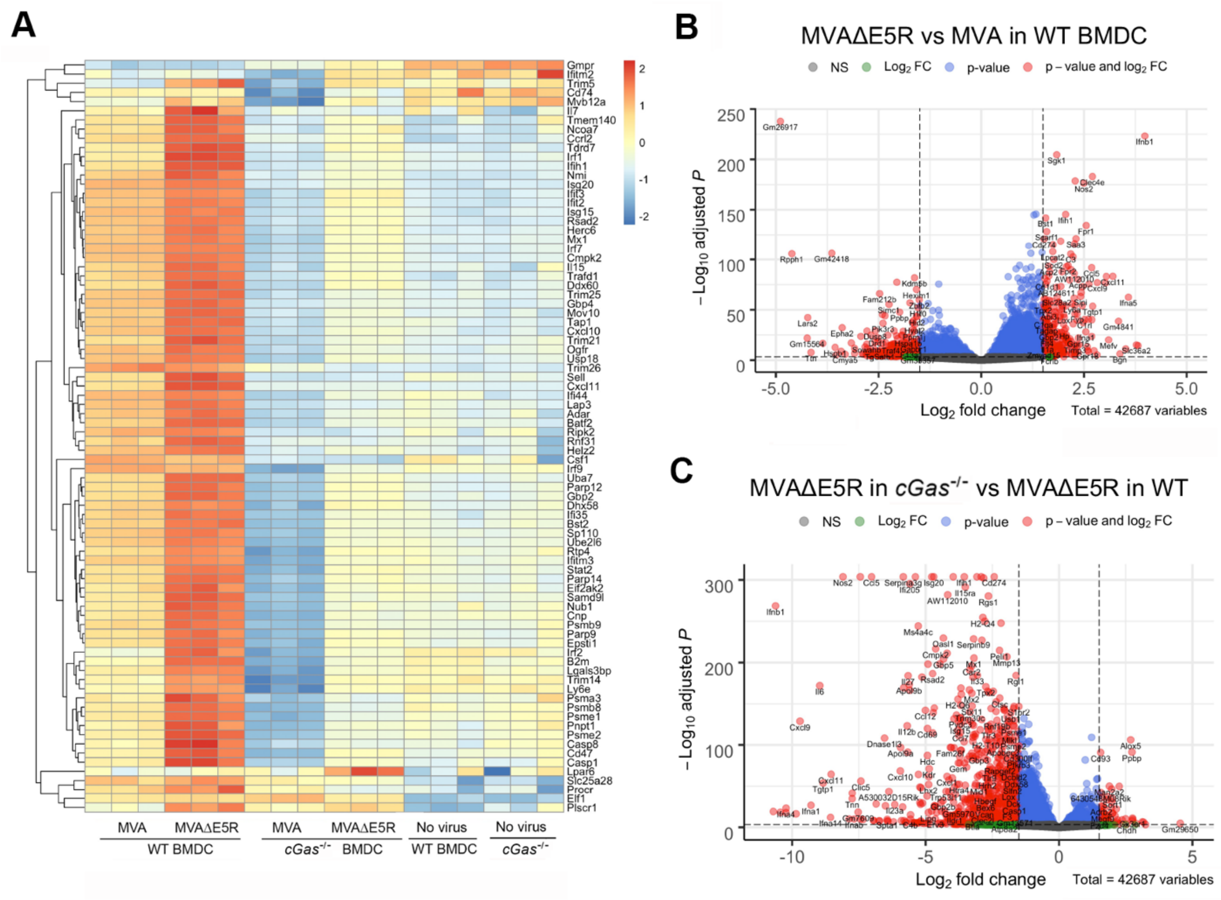


**Figure S4 related to Figure 6. MVAΔE5R-OVA promotes DC maturation and antigen-specific CD8$^{+}$ T cells activation.**

(A) A heatmap showing differentially expressed genes (DEGs) in WT or *cGas*$^{-/-}$ BMDCs infection with MVA vs. MVAΔE5R.

(B) A volcano plot showing DEGs in WT BMDCs infected with MVAΔE5R vs. MVA.

(C) A volcano plot showing DEGs in MVAΔE5R-infected WT vs. *cGas*$^{-/-}$ BMDCs.

#### Supplementary Table 1

List of primers used in this paper for qRT-PCR.

| **Primer sequence** | SOURCE |
|---|---|
| qPCR *Ifnb* For: TGGAGATGACGGAGAAGATG | Integrated DNA technologies IDT |
| qPCR *Ifnb* Rev: TTGGATGGCAAAGGCAGT | |
| qPCR *Ccl4* For: GCCCTCTCTCTCCTCTTGCT | |
| qPCR *Ccl4* Rev: CTGGTCTCATAGTAATCCATC | |
| qPCR *Gapdh* For: AGGTCGGTGTGAACGGATTTG | |
| qPCR *Gapdh* Rev: TGTAGACCATGTAGTTGAGGTCA | |
| qPCR *Ccl5* For: GCCCACGTCAAGGAGTATTTCTA | |
| qPCR *Ccl5* Rev: ACACACTTGGCGGTTCCTTC | |
| qPCR *Ifna* For: TCTGATGCAGCAGGTGGG | |
| qPCR *Ifna* Rev: AGGGCTCTCCAGACTTCTGCTCTG | |
| qPCR *A34* For: GGCATAGGAACATTTCTGCATTAC | |
| qPCR *A34* Rev: TACGACACTGATAAACCGCATT | |
| qPCR *A27* For: CCGTCCAGTCTGAACATCAAT | |
| qPCR *A27* Rev: GTGTTGTAAACGCAACGATGAA | |
